# Fibrillar adhesion dynamics govern the timescales of nuclear mechano-response via the vimentin cytoskeleton

**DOI:** 10.1101/2023.11.08.566191

**Authors:** Amy E.M. Beedle, Anuja Jaganathan, Aina Albajar-Sigalés, F. Max Yavitt, Kaustav Bera, Ion Andreu, Ignasi Granero-Moya, Dobryna Zalvidea, Zanetta Kechagia, Gerhard Wiche, Xavier Trepat, Johanna Ivaska, Kristi S. Anseth, Vivek B. Shenoy, Pere Roca-Cusachs

## Abstract

The cell nucleus is continuously exposed to external signals, of both chemical and mechanical nature. To ensure proper cellular response, cells need to regulate not only the transmission of these signals, but also their timing and duration. Such timescale regulation is well described for fluctuating chemical signals, but if and how it applies to mechanical signals reaching the nucleus is still unknown. Here we demonstrate that the formation of fibrillar adhesions locks the nucleus in a mechanically deformed conformation, setting the mechanical response timescale to that of fibrillar adhesion remodelling (∼1 hour). This process encompasses both mechanical deformation and associated mechanotransduction (such as via YAP), in response to both increased and decreased mechanical stimulation. The underlying mechanism is the anchoring of the vimentin cytoskeleton to fibrillar adhesions and the extracellular matrix through plectin 1f, which maintains nuclear deformation. Our results reveal a mechanism to regulate the timescale of mechanical adaptation, effectively setting a low pass filter to mechanotransduction.

## Introduction

Mechanical force is a fundamental regulator of cellular behaviour, driving changes in protein conformation and localisation, gene expression and cell function. The inability of a cell to correctly sense force underpins a number of pathologies, including fibrosis and cancer^1,2^. Mechanistically, when a cell receives a mechanical stimulus (such as force application or increased substrate rigidity) from the extracellular matrix (ECM), this triggers a highly coordinated chain of events that propagates the signal across the cytoplasm to the nucleus. This mechanotransduction process includes the growth and maturation of the focal adhesion complexes at the cell surface, and the formation and organization of actin stress fibres, which connect to and mechanically deform the nucleus^3^. In turn, nuclear deformation has a plethora of effects, including among others chromatin reorganization^4,5^, signalling at the nuclear envelope^6,7^, and altered nucleo-cytoplasmic transport dynamics, leading to the nuclear accumulation of mechanosensitive transcription factors^3,8^. Whereas these steps describe cell responses to increased force transmission, cells in physiological conditions are exposed to a dynamically changing environment where forces can also decrease. However, the molecular mechanisms and timescales that govern the reversibility of mechanotransduction are largely unknown.

A potential structure that could govern this reversibility, and its timescales, is the Extracellular Matrix (ECM) and its remodelling, which has recently been shown to store information of past cellular behaviour. Indeed, fibronectin deposition guides cell migration by generating a physiochemical cue that provides spatial memory^9^, and collagen remodelling promotes the invasion from a mechanically stiff to a soft environment via energy minimization^10^. ECM deposition and remodelling is also a defining feature of cells in a high rigidity environment^11^. This remodelling occurs concomitantly with the formation of fibrillar adhesions, which are long-lived integrin-α_5_β_1_ rich adhesions that colocalise with fibronectin fibrils. Fibrillar adhesions mature from focal adhesions as they are pulled by actin fibres and get progressively enriched with the protein tensin. From this evidence, it is tempting to hypothesize that ECM remodelling, and fibrillar adhesions, can regulate the dynamics of cell adaptation to a loss of forces.

Here we show that actomyosin contractility is required to initiate, but not sustain, nuclear deformation and subsequent mechanosignalling. Instead, nuclear deformation can be sustained simply through the anchoring of the vimentin cytoskeleton to the ECM through fibrillar adhesions. Upon loss of mechanical forces, this ECM-vimentin coupling delays mechano-adaptation by maintaining nuclear deformation and the nuclear localisation of mechanosensitive transcription factors. Furthermore, this ECM-vimentin connection also buffers high mechanical loads, protecting the nucleus from deformation and damage. Taken together, we unveil a mechanism by which fibrillar adhesions act as a low-pass filter for mechanical stimulation, setting the timescale of response to that of fibrillar adhesion remodelling (∼1 hour).

## Results

### Nuclear YAP is maintained upon loss of contractility in the presence of fibrillar adhesions

We sought to study the mechanisms regulating the re-localisation of mechanosensitive transcription factors upon loss of active contractile forces. We first seeded telomerase immortalized foreskin fibroblasts (TIFF) on fibronectin-coated glass coverslips for 4 hours to obtain a highly mechanically active phenotype, with the mechanosensitive transcription factor YAP localised to the nucleus. We then treated cells for 30 minutes with different pharmacological inhibitors that interfere with the actomyosin cytoskeleton. We found that there was no change in the Nuclear/Cytoplasmic (N/C) ratio of YAP upon treatment with blebbistatin (bleb, myosin inhibitor) or cytochalasinD (cytoD, actin inhibitor), whereas treatment with Y-27632 (Y-27, ROCK inhibitor) or latrunculinA (latA, actin inhibitor) triggered a decrease in nuclear localised YAP (Fig. 1a,b). Using traction force microscopy, we verified that all treatments dramatically decreased active cellular forces (fig. 1c,d), and therefore the maintenance of nuclear YAP localization is not explained by sustained cellular force generation. To understand the underlying mechanisms, our first approach was to study known markers of mechanotransduction, such as actin stress fibre organisation and focal adhesion length. Surprisingly, the change in localisation of YAP upon different pharmacological treatments did not correlate with these parameters (SI fig. 1). Instead, using a epitope-specific integrin-α5 antibody to mark fibrillar adhesions^12,13^ (Snaka51), we observed that in conditions where YAP remained nuclear (blebbistatin and cytochalasinD) the fibrillar adhesions were present, and conditions with loss of nuclear YAP (Y-27632 and LatrunculinA) correlated with loss of fibrillar adhesions (Fig. 1e,f). To further investigate this relationship, we varied the pharmacological treatment time (30 min – 2 hr) and found a good correlation (R^2^ = 0.7977) between the presence of fibrillar adhesions and the N/C ratio of YAP, where the loss of fibrils correlates with a loss of nuclear YAP (Fig. 1g). To elucidate whether this phenomenon was specific to the YAP signalling pathway, we performed analogous experiments with other known mechanosensitive transcription factors twist^14^ and snail^15^. These experiments recapitulated the findings with YAP, indicating that this is a general phenomenon regulating the cytoplasmic relocalisation of mechanosensitive molecules upon loss of active cellular forces (SI fig. 1).

**Figure 1.**
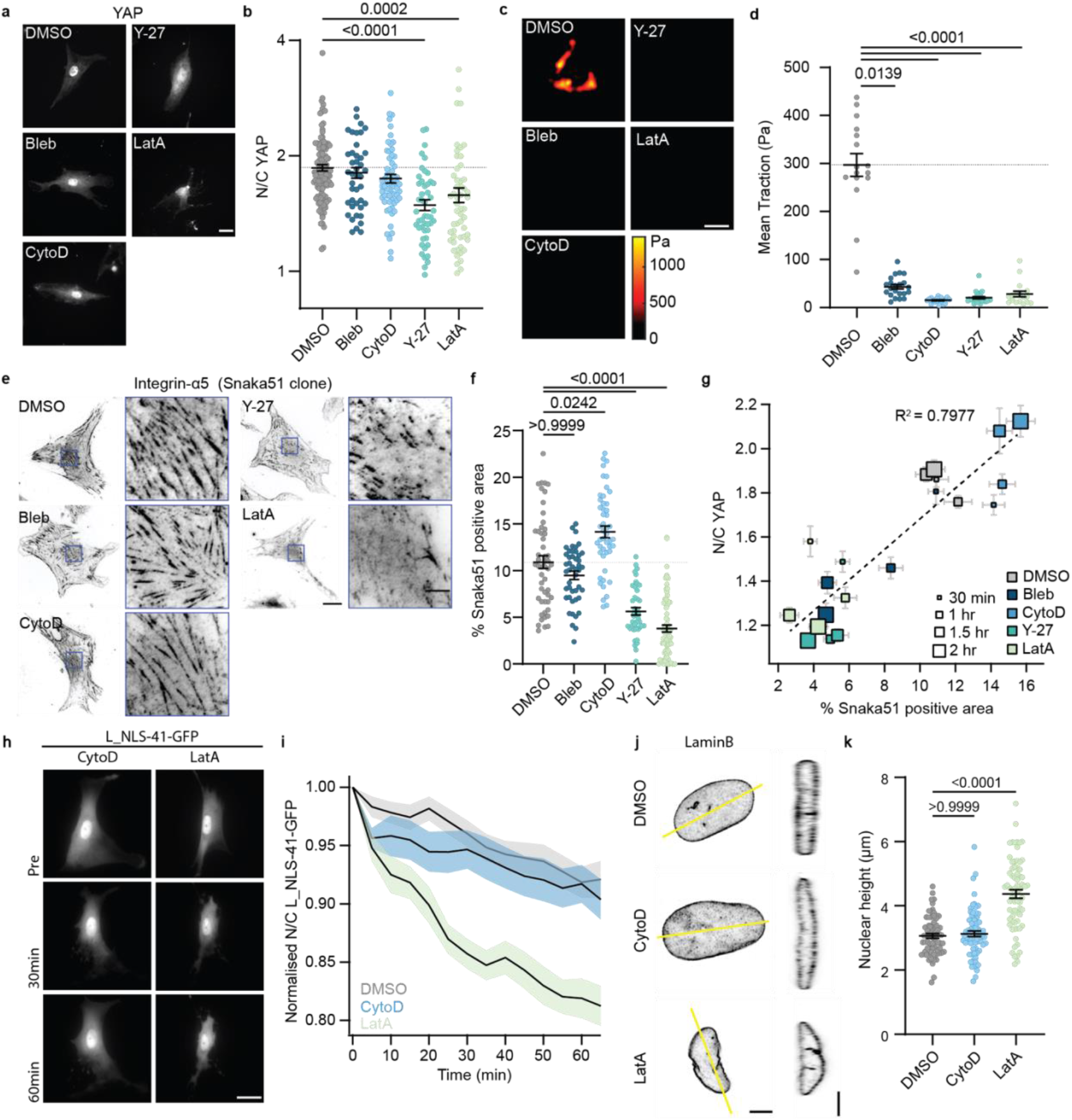
Mechanosensitive molecules remain in the nucleus upon loss of contractility when fibrillar adhesions are present. **a.** Example images of YAP stained cells after 30min of indicated pharmacological treatment; scale bar 25μm. **b.** Quantification of N/C YAP ratio after 30min pharmacological treatment. (n=121/42/60/48/57 cells for DMSO/bleb/cytoD/Y-27/latA from at least 3 independent experiments; Kruskal-wallis analysis of variance (ANOVA) with Dunn’s multiple comparison test). **c.** Example colour maps of traction forces after 30min pharmacological treatment; scale bar 50μm. **d.** Quantification of cell tractions on 15kPa polyacrylamide gels. (n=16/22/21/26/17 cells for DMSO/bleb/cytoD/Y-27/latA from 2 independent experiments; Kruskal-wallis ANOVA with Dunn’s multiple comparison test). **e.** Examples of integrin-α_5_ snaka51 clone stained cells after 30min of the indicated pharmacological treatment; scale bar 25μm / zoom 5μm. **f.** Quantification of the percentage area under the nucleus occupied by fibrillar adhesions. (n=52/46/48/45/76 cells for DMSO/bleb/cytoD/Y-27/latA from at least 3 independent experiments; Kruskal-wallis ANOVA with Dunn’s multiple comparison test). **g.** Correlation between N/C YAP ratio and percentage area of fibrillar adhesions under the nucleus for different drug treatments (colour coded) with different incubation times (size coded). (At least 3 independent experiments with a minimum of 28 cells per condition). **h.** Example images of mechano-reporter L_NLS-41-GFP transfected cells prior to and during cytoD or latA treatment; scale bar 25μm. **i.** Quantification of change in N/C L_NLS-41-GFP ratio with time (normalized to the pre-treatment ratio) for cells upon with indicated pharmacological treatment. (n=40/33/30 cells for DMSO/cytoD/latA from 3 independent experiments). **j.** Examples of LaminB stained nuclei after 1 hour of the indicated drug treatment. The yellow line signifies the position of the Z-plane re-slice; scale bar 5µm. **k.** Quantification of the nuclear height after 1hr pharmacological treatment. (n=69/70/70 nuclei for DMSO/cytoD/latA from 5 independent experiments; Kruskal-wallis ANOVA with Dunn’s multiple comparison test)

To decouple the contributions of mechanosensitivity and biochemical signalling regulating the change in molecular localisation, we performed experiments using a previously developed mechano-reporter (L_NLS-41-GFP). This reporter functions independently of chemical signalling, and instead responds to mechanically-induced changes in facilitated and passive nucleocytoplasmic diffusion such that it localises to the nucleus in higher rigidity environments^8^ (SI fig. 1). We transfected cells with the mechano-reporter and treated them with either cytoD or latA, both targeting the actin cytoskeleton but with differential effects on the fibrillar adhesions. Upon treatment with cytoD, which does not disrupt fibrillar adhesions, the dynamics of the reporter was indistinguishable from the control DMSO treatment (Fig. 1h,i). By contrast, treating cells with latA, which disrupts fibrillar adhesions, caused a rapid loss of the sensor from the nucleus (Fig. 1h,i). Given that the localisation of mechanosensitive molecules such as YAP, snail or twist1 is caused by force transmission to the nucleus, and subsequent nuclear deformation and flattening^3,8^, we thus sought to understand whether these differences in nuclear localisation are associated with changes in nuclear morphology. After 1 hour cytoD treatment, the nuclear height was not significantly different from the control DMSO condition (Fig 1j,k) despite the total removal of actin stress fibres spanning the nucleus (SI fig. 1). However, in cells treated for 1 hour with latA, the nuclei undergo a significant increase in nuclear height (Fig. 1j,k). Taken together, these results suggest that fibrillar adhesions maintain nuclear morphology and delay the relocalisation of mechanosensitive transcription factors from the nucleus upon loss of contractile forces.

### Inhibiting fibrillar adhesion formation leads to rapid loss of mechanosensitive molecules from the nucleus and altered nuclear morphology upon loss of contractility

To test the hypothesis that fibrillar adhesions alter cell response to loss of contractile forces, we impeded the formation of fibrillar adhesions by inhibiting the remodelling of fibronectin. As a first approach, we used the PUR4 (also known as FUD) peptide that binds with high affinity to N-terminus of fibronectin and is a potent inhibitor of the assembly of fibronectin into fibrils^16–18^ (SI fig. 2). This effectively prevented the formation of fibrillar adhesions (Fig. 2a), but did not alter key aspects of cellular mechanotransduction, including focal adhesion growth, and the nuclear localisation of YAP (SI Fig. 2, Fig. 2c,d). To understand the contribution of fibrillar adhesions upon loss of contractile forces, we exposed cells to either PUR4 or a control scrambled peptide and treated them with blebbistatin or cytochalasinD. As expected, for the control conditions these treatments did not trigger a decrease in the N/C ratio of YAP (Fig. 2c,d). However, in the cells lacking fibrillar adhesions, this pharmacological treatment triggered a significant decrease in the N/C YAP ratio (Fig. 2c,d). The same results were obtained when we tracked the mechano-reporter L_NLS-41 over time in cytoD treated cells, showing the generality of the results beyond YAP (Fig. 2e,f). This change in localisation was also associated with a change in nuclear morphology. When the fibrillar adhesions are present (exposure to control peptide), nuclear height was not affected by cytoD (Fig. 2g,h). By contrast, when fibrillar adhesion formation was inhibited (PUR4 exposure), the cytoD treatment increased nuclear height (Fig. 2g,h).

**Figure 2.**
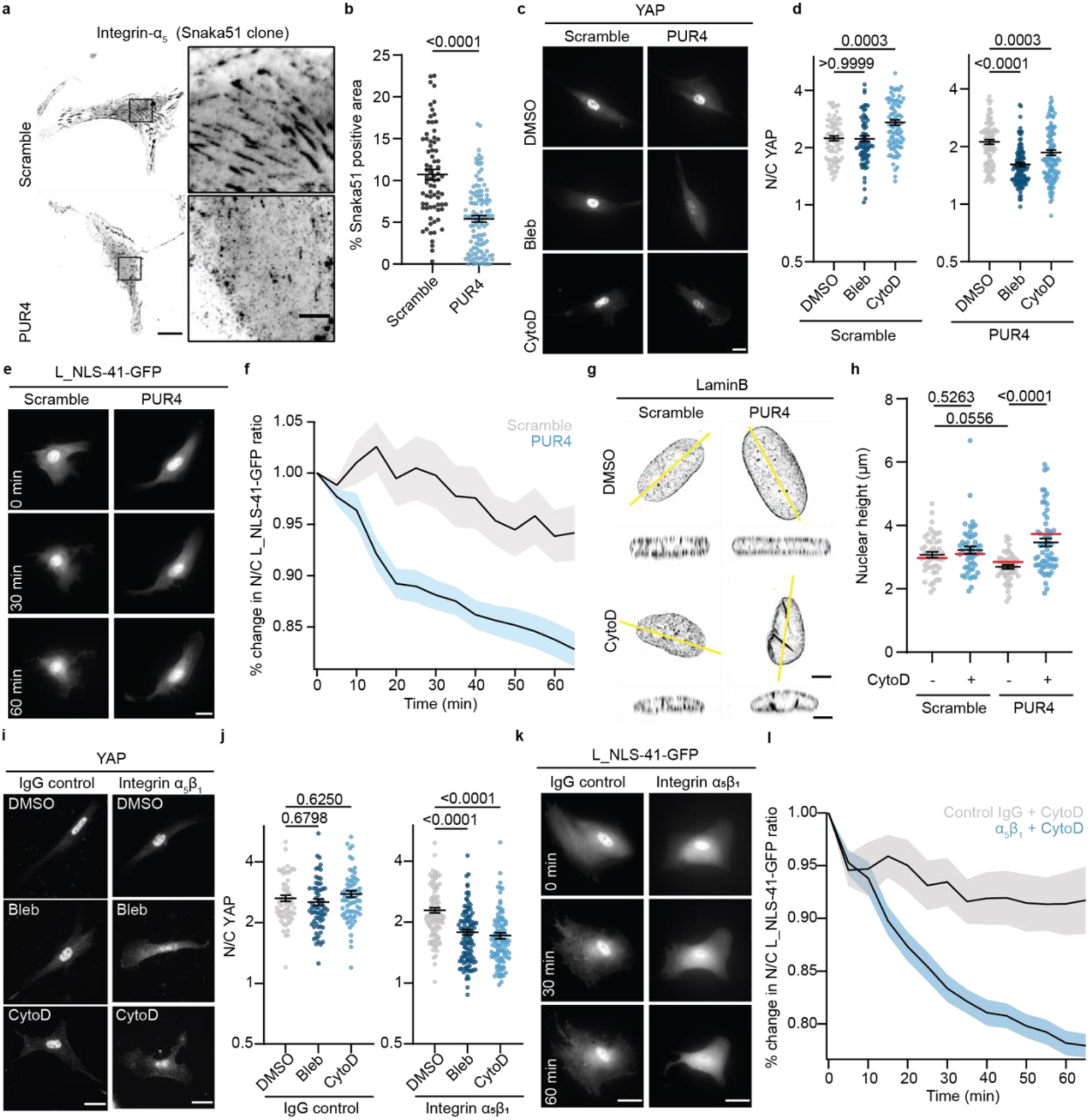
Inhibition of fibrillar adhesions leads to rapid loss of mechanosensitive molecules when contractility is inhibited. Fibrillar adhesions maintain nuclear morphology in absence of actin cap. **a.** Example images of integrin-α_5_, clone snaka51 stained cells in scramble or PUR4 peptide seeded for 4 hours in presence of 500nM peptide. Scale bar 20µm, zoom 5µm. **b.** Percentage area under the nucleus occupied by fibrillar adhesions for scramble and PUR4 peptide. (n=74/109 cells for scramble/PUR4 from 4 independent experiments; unpaired t-test). **c.** Example images of YAP stained cells treated for 30min with indicated pharmacological treatment for scramble and PUR4 peptide. Scale bar 20µm. **d.** Quantification of N/C YAP ratio for cells in scramble or PUR4 peptide conditions treated with 30min pharmacological treatment. (Scramble – n=80/74/82 cells for DMSO/bleb/cytoD. PUR4 – n=93/118/110 cells for DMSO/bleb/cytoD from at least 3 independent experiments; Kruskal-wallis ANOVA with Dunn’s multiple comparison test). **e.** Example images of mechano-reporter L_NLS-41-GFP transfected cells in scramble or PUR4 peptide at different time points are treatment with cytoD. Scale bar 20µm. **f.** Quantification of change in N/C L_NLS-41-GFP ratio (normalized to the pre-cytoD treatment ratio) for scramble (grey) and PUR4 (blue) peptide conditions treated with cytoD. Scramble n=36, PUR4 n=44 cells from 3 independent experiments. **g.** Example images of LaminB stained nuclei for scramble and PUR4 cells treated with DMSO control or CytoD. Yellow line indicates the position of the re-slice; sale bar 5µm. **h.** Quantification of nuclear height for scramble and PUR4 cells in the absence and presence of cytoD. (Scramble – n=46/49 nuclei for DMSO/cytoD. PUR4 – n=48/60 nuclei for DMSO/cytoD from 3 independent experiments; two-way ANOVA with Tukey’s multiple comparisons test). The red bars represent the values of nuclear height obtained from the computational model. **i.** Example images of YAP stained cells for cells blocked with IgG control or Integrin α_5_β_1_ antibody and treated with indicated pharmacological treatment for 30 minutes. Scale bar 20µm. **j.** Quantification of N/C YAP ratio for cells blocked with IgG control or Integrin α_5_β_1_ and subjected to 30min pharmacological treatment. (IgG – n=60/65/67 cells for DMSO/bleb/cytoD. Integrin α_5_β_1_ – n=103/109/109 cells for DMSO/bleb/cytoD from at least 4 independent experiments; Kruskal-wallis ANOVA with Dunn’s multiple comparison test). **k.** Example images of mechano-reporter L_NLS-41-GFP transfected cells treated with IgG control or integrin α_5_β_1_ antibody at different timepoints after treatment with cytoD. Scale bar 20µm. **l.** Quantification of change in N/C L_NLS-41-GFP ratio (normalized to the pre-cytoD treatment ratio) for control IgG blocked cells treated with cytoD (grey) and integrin α5β1 blocked cells treated with cytoD (blue). Solid line represents the average of all trajectories and shaded area is the standard error. IgG n=21 cells, Integrin α5β1 n=25 cells from 5 independent experiments.

To further confirm the role of fibrillar adhesions, we interfered with their formation with five alternative methods. First, we crosslinked fibronectin with glutaraldehyde prior to cell seeding, which prevents its remodelling and subsequent fibrillar adhesion formation^19^. This led to the same trends in YAP and nuclear height as the PUR4 peptide (SI Fig 2). Second, we used a blocking antibody against α_5_β_1_ integrins, through which fibrillar adhesions attach to fibronectin^20^. Cells with blocked α_5_β_1_ lacked fibrillar adhesions (SI Fig. 2) but exhibited nuclear localised YAP (Fig. 2i,k). Upon treatment with blebbistatin or cytochalasinD, cells treated with α_5_β_1_ antibody, but not with a control antibody, rapidly lost the nuclear localisation of both YAP (Fig. 2i,j) and the mechano-reporter (Fig. 2k,l). This was accompanied by a change in nuclear morphology (SI Fig. 2). Third, we compared cells seeded on high rigidity gels (30 kPa), where fibrillar adhesions were formed and YAP was nuclear, to intermediate rigidity gels (5 kPa), where YAP was already nuclear but fibrillar adhesions were smaller as described previously^12^ (SI Fig. 3). Upon treatment with cytoD, the cells on the higher rigidity gels maintained the N/C YAP ratio, whereas there was a significant reduction in the cells seeded on 5 kPa gels (SI fig. 3). Fourth, we applied bleb or cytoD to MCF10A mammary epithelial cells, which do not form fibrillar adhesions, and observed a significant reduction in N/C YAP ratio after 30 min (SI fig. 3).

**Figure 3:**
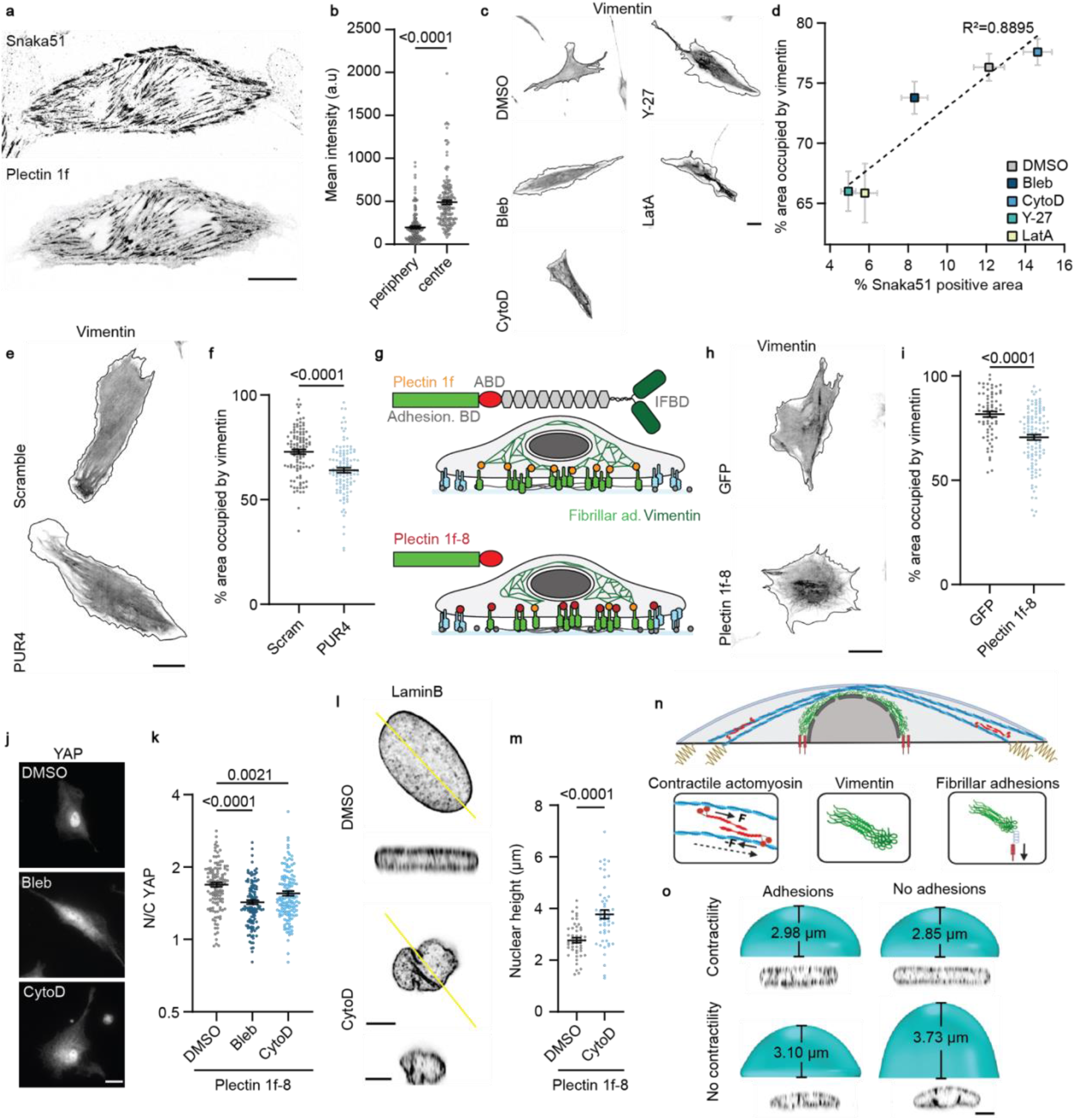
Fibrillar adhesions anchor the vimentin network via plectin 1f and maintain nuclear morphology in absence of active forces. **a.** Example of cell transfected with plectin 1f-GFP and stained with snaka51 antibody. Scale bar 20µm. **b.** The mean intensity of plectin 1f at snaka51 adhesions close to the cell periphery and in the central region of the cell. (n=160 adhesions from 32 cells from 3 independent experiments; Mann-Whitney test). **c.** Example images of vimentin morphology in cells treated with indicated pharmacological treatment for 1 hour. Black line indicates the cell outline. Scale bar 20 µm. **d.** The percentage area of the cell occupied for vimentin vs the percentage area under the nucleus occupied by fibrillar adhesions for 1 hour pharmacological treatment (colour coded). (Vimentin – n=59/59/59/57/47 cells for DMSO/bleb/cytoD/Y-27/latA from 3 independent experiments. Fibrillar adhesions – n=123/42/60/48/57 cells for DMSO/bleb/cytoD/Y-27/latA from at least 3 independent experiments). **e.** Example of vimentin stained cells in the presence of the scramble or PUR4 peptide. Black line indicates the cell periphery. Scale bar 25µm. **f.** The percentage area of the cell occupied by vimentin in cells cultured with scrambled or PUR4 peptide. (n=108/112 cells for scramble/PUR4 from 7 independent experiments; Mann-Whitney test). **g.** Schematic representation of the full length plectin 1f and the truncated plectin 1f-8 lacking the intermediate filament binding domain. **h.** GFP and plectin 1f-8 transfected cells stained for vimentin. Solid black line indicates the perimeter of the cell. Scale bar 25 µm. **i.** Quantification of the percentage area of the cell occupied by the vimentin network in GPF and plectin 1f-8 transfected cells. (n=67/112 cells for GFP/ Plectin 1f-8 from at least 5 independent experiments; Mann-Whitney test). **j.** Example of YAP staining for plectin 1f-8-GFP expressing cells treated for 30min with the indicated pharmacological treatment. Scale bar 20 µm. **k.** Quantification of N/C YAP ratio of plectin 1f-8 transfected cells after 30min pharmacological treatment. (n=120/120/132 cells for DMSO/bleb/cytoD from 5 independent experiments; Kruskal-wallis ANOVA with Dunn’s multiple comparison test). **l.** Images of LaminB stained plectin 1f-8 transfected cells treated with DMSO and cytoD. Scale bar 5 µm. **m.** Quantification of nuclear height of plectin 1f-8 transfected cells treated with DMSO or cytoD. (n=48/52 nuclei for DMSO/cytoD from 3 independent experiments; unpaired two-tailed t-test). **n.** Scheme showing main computational model components. This includes actomyosin contractility driven by myosin motors, and a vimentin network around the nucleus anchored to the substrate via fibrillar adhesions. Fibrillar adhesions are adhesive and withstand cell/ECM forces in a dissipative way. **o.** Model predictions for nuclear height with anchored/unanchored vimentin (adhesions/no adhesions) and in the presence/absence of contractility. Experimental data from fig. 2g are shown below model predictions for comparison.

Finally, given that fibrillar adhesions form upon the maturation of focal adhesions, there should be a time-dependent effect linked to fibrillar adhesion formation. To test this, we first probed the timescales of focal adhesion formation, fibrillar adhesion formation, and YAP localization and observed that after 2 hours of seeding focal adhesions are formed and YAP is localised to the nucleus, but the fibrillar adhesions were not fully mature (SI fig 3). We subsequently inhibited contractility in cells seeded for 2 hours and observed a significant change in YAP localisation at short timescales (SI fig 3). This is in contrast to the lack of effect observed 4 hours after cell seeding, when fibrillar adhesions are fully formed (Fig. 1b). Altogether, these results demonstrate that fibrillar adhesions maintain the localisation of mechanosensitive molecules in the absence of contractility by sustaining a deformed, flat nuclear morphology independently of the actin cytoskeletal network.

### Fibrillar adhesions anchor the vimentin network via plectin 1f

To explore the underlying mechanism by which fibrillar adhesions regulate nuclear morphology and mechanosensitive molecular localisation upon loss of contractile forces, we hypothesized that there may be a contribution from cytoskeletal components. Given that YAP remained nuclear in the absence of an actin network (cytoD treatment, Fig. 1), we turned our attention to other cytoskeletal networks. In particular, fibrillar adhesions serve as docking sites for vimentin via the cytolinker protein plectin isoform 1f^21^ and thus fibrillar adhesions may regulate the organisation of the vimentin intermediate filament network. To assess this, we first transfected cells with plectin 1f-GFP and performed stainings against the fibrillar adhesion marker snaka51 (Fig. 3a). Confirming the presence of plectin 1f in fibrillar adhesions, plectin 1f and snaka51 colocalized in the central region of cells (where snaka51 marks fibrillar adhesions) and not at the periphery (where snaka51 marks focal adhesions, Fig. 3b). Then, we assessed the organization of the vimentin network upon applying the different perturbations used in figures 1 and 2. In response to pharmacological treatments, we found that conditions with fibrillar adhesion loss also exhibited a collapsed vimentin network, measured as a reduction in the percentage area of the cell occupied by vimentin (Fig. 3c,d), with a very high correlation between both parameters (R^2^=0.8895). Upon blocking fibronectin remodelling with either the PUR4 peptide (Fig. 3e,f) or glutaraldehyde (SI Fig. 4), the cell area occupied by vimentin decreased. Finally, vimentin spreading was also increased along fibrillar adhesions in response to substrate stiffness (SI Fig. 4). Thus, the ability of vimentin to spread and form a structured cytoskeleton is determined by the cells ability to remodel fibronectin, and the subsequent formation of fibrillar adhesions.

**Figure 4:**
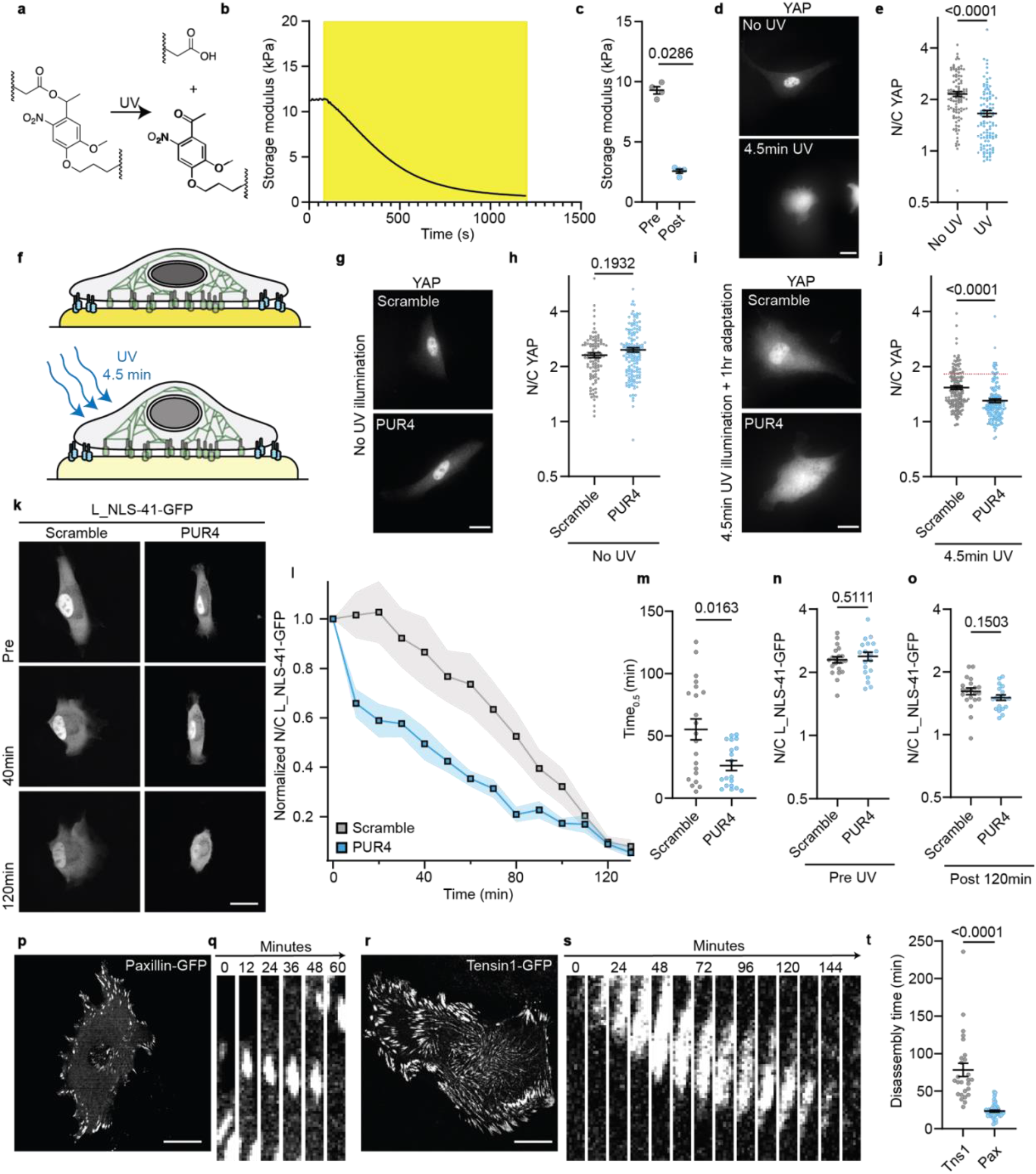
The ECM-vimentin coupling delays mechanoadaptation to a soft environment. **a.** Chemical structure of the photodegradable compound. **b.** Storage modulus of the photodegradable gel, which softens upon illumination with UV light (yellow region, 365 nm, 10 mW cm^-2^). Trace representative of 3 gels. **c.** Quantification of the gel storage modulus before UV exposure and after 5min UV exposure (n=4 gels, *p* = 0.0286 from Mann-Whitney test). **d.** Examples of YAP staining in cells seeded on gels that were not softened, or softened with 4.5min of UV illumination prior to cell seeding; scale bar 20 µm. **e.** Quantification of N/C YAP ratio in cells seeded on photodegradable gels without UV exposure, or with UV exposure prior to cell seeding. (n=98/107 cells for NoUV/UV from 3 independent experiments; Mann-Whitney test). **f.** Schematic representation of experimental set-up, whereby cells are seeded on photodegradable gels (with scramble or PUR4 peptide) for 4hours. The gel is then softened with 4.5min of UV illumination. **g.** Examples of YAP staining of cells seeded on photodegradable gels without UV exposure in scramble and PUR4 condition; scale bar 20 µm. **h.** Quantification of N/C YAP ratio for cells in scramble or PUR4 peptide seeded on gels without UV softening. (n=100/141 for scramble/PUR4 from 3 independent experiments; Mann-Whitney test). **i.** Examples of YAP staining of cells on photodegradable gels 1hour after a 4.5min UV exposure for scramble and PUR4 condition; scale bar 20 µm. **j.** Quantification of N/C YAP ratio for cells in scramble of PUR4 peptide 1 hour after UV induced gel softening. The red line corresponds to the mean N/C YAP ratio for cells seeded on non-degradable polyacrylamide gels and subjected to the same UV conditions; additional details in SI fig. 4. (n=144/170 cells for scramble/PUR4 from 3 independent experiments; Mann-Whitney test). **k.** Examples of mechano-reporter L_NLS-41-GFP transfected cells on photodegradable gels pre-, after 40min-, and 120min-UV illumination for the scramble and PUR4 condition; scale bar 20 µm. **l.** Quantification of N/C L_NLS-41-GFP ratio with time (normalized to the initial and final ratio) for cells on photodegradable gels upon exposure to 4.5min UV illumination for cells in scramble (grey) or PUR4 (blue) peptide. (n=21/19 cells for scramble/PUR from 3 independent experiments). **m.** Quantification of the time taken for the L_NLS-41-GFP ratio to fall below 50% of the initial value for scramble and PUR4. n=21/19 cells for scramble/PUR from 3 independent experiments; Mann-Whitney test) **n.** Quantification of N/C L_NLS-41-GFP ratio for cells seeded on photodegradable gels prior to UV illumination. (n=21/19 cells for scramble/PUR from 3 independent experiments; Unpaired two-tail t-test). **o.** Quantification of N/C L_NLS-41-GFP ratio 120min after the UV exposure for scramble and PUR4 condition. (n=21/19 cells for scramble/PUR from 3 independent experiments; Unpaired two-tail t-test). **p.** Example image of cell expressing paxillin-GFP seeded on glass; scale bar 20 µm. **q.** Images of an example paxillin-GFP adhesion at 12 minute intervals; scale bar is 1 µm. **r.** Example image of cell expressing tensin1-GFP seeded on glass; scale bar 20 µm. **s.** Images of an example tensin1-GFP adhesion at 12 minute intervals; scale bar is 1 µm. **t.** Quantification of adhesion disassembly time of paxillin and tensin1 adhesions. (Paxillin n=47 adhesions from 11 cells, 5 independent experiments and tensin1 n=26 adhesions from 10 cells, 4 independent experiments; Mann-Whitney test).

Given that the fibrillar adhesion – vimentin connection is mediated by plectin 1f, breaking this connection should lead to vimentin network collapse regardless of the presence of fibrillar adhesions. To investigate this, we transfected cells with a truncated version of plectin 1f which contains the fibrillar adhesion-binding N-terminal domain but lacks the intermediate filament-binding C-terminal domain^21^ (plectin 1f-8) (Fig. 3g). Cells overexpressing plectin 1f-8-GFP formed fibrillar adhesions where plectin 1f-8-GFP localized (SI. Fig 4), but exhibited reduced vimentin spreading as compared to cells transfected with GFP alone (Fig. 3h,i). Thus, plectin 1f-8 functions as a dominant-negative, likely by displacing some of the endogenous plectin 1f and reducing the connectivity between the fibrillar adhesions and vimentin. Consistently, plectin 1f-8 overexpression did not affect N/C YAP ratios in control untreated cells (SI fig. 4), but it abolished their ability to retain nuclear YAP upon contractility inhibition (Fig. 3j,k and SI Fig. 4). Furthermore, upon treatment with cytoD which destroys the actin cap, cells expressing GFP-alone maintained their nuclear morphology (SI. Fig 4), but in plectin 1f-8 expressing cells the nuclear height significantly increased (Fig. 3l,m). Thus, a vimentin network properly anchored by plectin 1f is able to sustain a deformed nuclear morphology in the absence of contractile forces and a compressive actin cap, maintaining the localisation of the mechanosensitive transcription factor YAP.

To further understand whether a simple vimentin cage around the nucleus could maintain nuclear shape, we generated a mechanical model considering the key elements involved. The model (see methods) considers a contractile actomyosin network anchored to the cell periphery via focal adhesions. A vimentin network spans the nucleus and is anchored to fibrillar adhesions via adhesive interactions. In conditions lacking fibrillar adhesions, this adhesive force is set to zero effectively decoupling vimentin from the extracellular environment. We first allowed the contractile actin network to form, leading to high compressive strains and a flattening of the nucleus (Fig. 3n). Regardless of whether vimentin is anchored, we observed a similar nuclear morphology that is consistent with the experimental observations. We subsequently removed contractility from the model and observed the effect on nuclear morphology. In the condition where vimentin is anchored to fibrillar adhesions, the adhesive force prevents any change in nuclear morphology. This is because once the fibrillar adhesions are engaged with the vimentin cage the interactions persist even when the contractility is abrogated. While contractility is not needed to sustain these bonds, it is essential to initiate their formation. By contrast, in the case where vimentin is not anchored, we observe a significant rounding of the nucleus (Fig. 3o and red bars in Fig. 2h). This is consistent with the experimental results, and demonstrates that vimentin anchoring to the substrate is sufficient to maintain a compressed nuclear morphology in the absence of active contractile forces.

### The ECM-vimentin coupling alters mechano-adaptation timescales

Thus far, we have shown that the connectivity between the fibrillar adhesions and the vimentin network delays the loss of mechanical signals upon inhibition of cellular contractility. Next, we asked whether this mechanism would also determine the timescales by which cells adapt from a high to low rigidity mechanical environment. Indeed, by maintaining a mechanically active phenotype, fibrillar adhesions could delay the timescales of adaptation from a stiff-to-soft conversion. As an initial experiment, we treated cells seeded on low rigidity polyacrylamide gels (1.5kPa) with Mn^2+^, which activated α_5_β_1_ integrins and initiated cell spreading, fibrillar adhesion formation, and nuclear localisation of the mechano-reporter, mimicking the mechanical activation of stiff gels (SI fig. 5). Upon removal of Mn^2+^, PUR4 treated cells (with blocked fibrillar adhesions) decreased nuclear area, and nuclear localization of the mechano-reporter, faster than cells treated with the scrambled peptide (SI Fig. 5).

**Figure 5:**
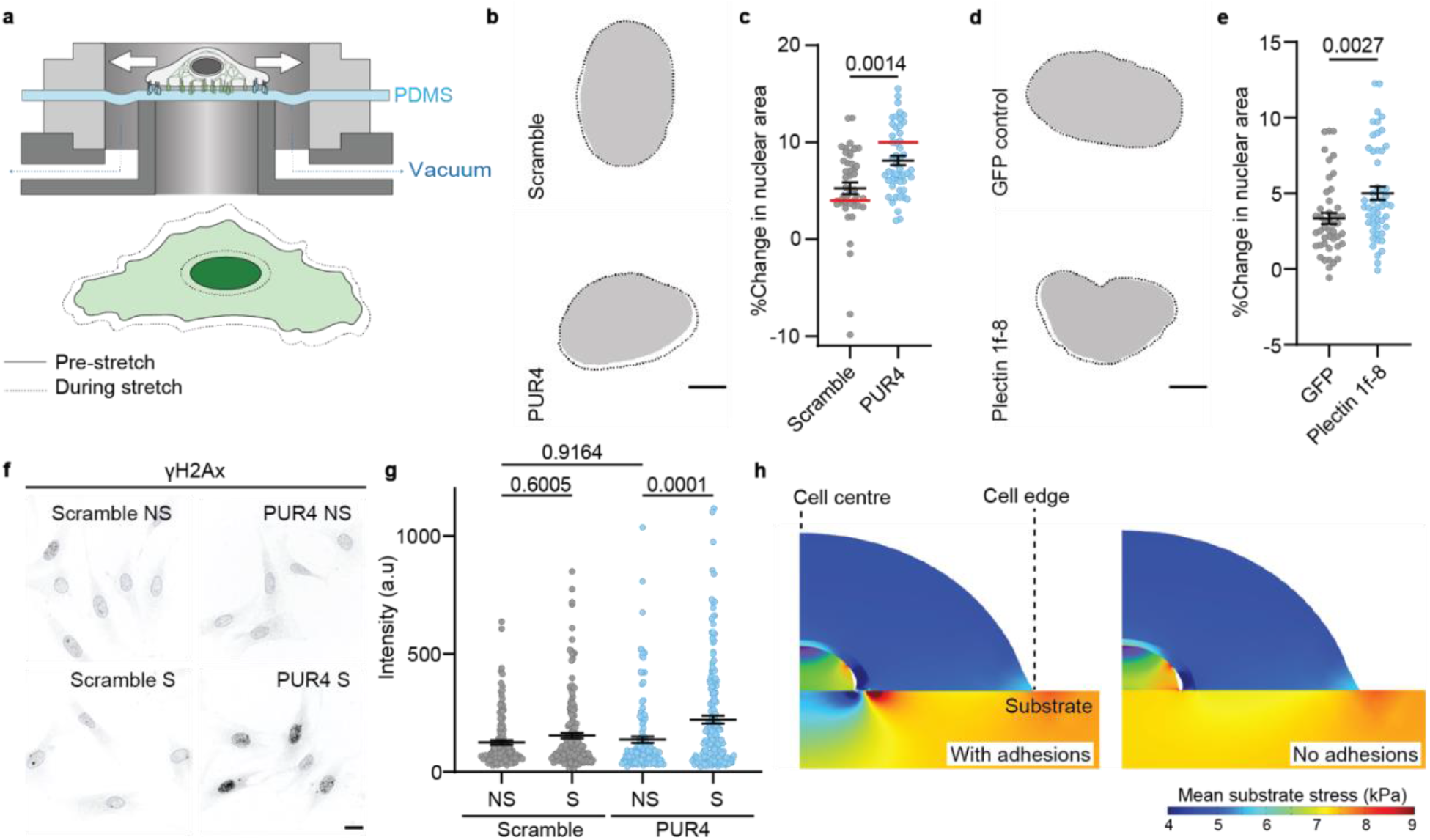
The ECM-vimentin coupling protects the nucleus from mechanical deformation and DNA damage. **a.** Schematic of the stretch device. An image of the membrane and the nucleus of is acquired before and during the application of a stretch. **b.** Example images of nuclear shape before (solid) and during (dashed line) stretch for cells in scramble or PUR4 peptide. Scale bar 5µm. **c.** Quantification of percentage change in nuclear area upon stretch for cells in scramble or PUR4 peptide. (n=47/50 cells for scramble/PUR4 from 3 independent experiments; Mann-Whitney two-tailed test). Red bars correspond to the percentage change in nuclear area obtained from the computational model. **d.** Example images of nuclear shape before (solid) and during (dashed line) stretch for cells transfected with GFP only (control) or plectin 1-8-GFP. Scale bar 5µm. **e.** Quantification of percentage change in nuclear area upon stretch for GFP or plectin 1f-8 transfected cells. (n=46/51 cells for GFP/plectin 1f-8 from 3 independent experiments; Mann-Whitney two-tailed test). **f.** Example images of γH2Ax stained cells cultured in scramble or PUR4 peptide that were not stretched (NS) or subjected to stretch (S). Scale bar 20µm. **g.** Quantification of γH2Ax nuclear intensity for cells in scramble or PUR4 peptide without stretch (NS) or after stretch (S). (n=132/174/139/175 cells for scramble-NS/Scramble-S/PUR4-NS/PUR4-S from 2 independent experiments; 2way ANOVA with Tukey’s multiple comparison test). **h.** Model predictions for stresses in the cell and the substrate during mechanical stretch for cells with an anchored vimentin network (adhesions), and cells lacking vimentin anchoring (no adhesions).

These experiments suggest a delay in mechano-adaptation, but do not fully mimic a change in substrate rigidity. To improve this, we fabricated stiff hydrogels that contain a photocleavable cross-linker that breaks upon illumination with UV light, triggering a softening of the hydrogel^22^ (Fig. 4a-c). The extent of softening can be regulated by the dose of light. We first verified that the gel softening affected cell mechanotransduction by seeding cells on unexposed gels (∼9.3 kPa in Young’s modulus), or gels softened for 4.5mins prior to cell seeding. As expected, N/C YAP ratio in cells seeded on the pre-softened gels (Fig. 4d,e) was significantly lower. We next utilized the photodegradable hydrogels to investigate whether fibrillar adhesions regulate how cells adapt to a change in the mechanical properties of the environment. We seeded cells on non-softened gels for >4 hours in the presence of the scramble or PUR4 peptide to control fibrillar adhesion formation and observed that in both conditions YAP was localized to the nucleus (Fig. 4f-h). We subsequently *in-situ* softened the gels with 4.5min UV illumination and waited for 1 hour to allow the cells to adapt to the new low rigidity environment. After 1 hour there was a reduction in the N/C YAP ratio for both conditions; however, the cells with fibrillar adhesions (scramble) had a significantly higher N/C YAP ratio compared to the cells lacking fibrillar adhesions (PUR4) (Fig. 4i,j), suggesting that fibrillar adhesions help to maintain a mechanically active phenotype and delays the adaptation timescales. Confirming that this effect is due to substrate mechanics rather than UV illumination, UV exposure to cells on non-degradable polyacrylamide gels of similar rigidity did not produce differences in N/C YAP between the two conditions (SI fig. 5).

To understand how fibrillar adhesions affect the timescales of mechanoadaptation, we performed *in-situ* softening experiments on cells transfected with the mechano-reporter (L_NLS-41-GFP) in the presence of the PUR4 or scrambled peptide (Fig. 4k-o). In the presence of the PUR4 peptide, the N/C ratio of the sensor began to decrease immediately upon softening (Fig. 4l). By contrast, control cells with fibrillar adhesions displayed a lag-time, where the N/C ratio of the sensor was unaffected by the gel softening for ∼30mins before decreasing. Correspondingly, the time required for the sensor N/C ratio to fall below half of the starting value (t_0.5_) was ∼52 minutes for control cells, compared to ∼17 minutes in the presence of the PUR4 peptide (Fig. 4m). We verified that the initial (pre-softened) and final (2 hour post softening) N/C sensor ratio was the same for both conditions (Fig. n,o). We therefore sought to understand whether these differences in cellular response timescales stem from differences in dynamics of the focal adhesions compared to the fibrillar adhesions. We analysed the disassembly timescales of focal adhesions (marked with paxillin-GFP, Fig. 4p,q) and fibrillar adhesions (marked with tenins1-GFP, Fig. 4r,s) and found a stark difference. While the focal adhesions disassembled within ∼20min, fibrillar adhesions required ∼60 mins (Fig. 4t), thereby closely matching the timescales of adaptation to soft substrates. Altogether, this demonstrates that the stable dynamics of fibrillar adhesions sets the timescales of cellular mechano-adaptation, thus delaying the relocalisation of mechanosensitive molecules upon softening of the mechanical environment.

### The ECM-vimentin coupling protects the nucleus from mechanical deformation

Thus far we have demonstrated that an anchored vimentin network sets the timescales for adaptation upon a loss of force. A more well-established role for the intermediate filament network^23,24^, and in particular vimentin^25,26^, is that it protects the nucleus from mechanical deformation and damage. However, this knowledge largely stems from studies comparing cells lacking a vimentin network to cells with an intact network. Our results raise the question of whether the vimentin network must merely be present or must be anchored to the ECM to effectively dissipate high mechanical loads. To address this question, we stretched cells by ∼10% and measured the corresponding change in cell and nuclear area (Fig. 5a). Cells treated with either PUR4 or the scrambled peptide increased their membrane area equally upon stretch (SI Fig. 6). Fibrillar adhesions (marked with tensin-GFP) were also stretched (SI Fig. 6). However, the nuclei of cells with the scrambled peptide (and therefore with fibrillar adhesions) increased their area to a smaller degree than cells exposed to PUR4 (Fig. 5b,c). The same trends were observed when blocking fibrillar adhesions with glutaraldehyde (SI Fig. 6), or with plectin 1f-8 overexpression (Fig 5d,e and SI fig 6). We therefore hypothesised that the anchoring of vimentin allows the vimentin network to stretch and dissipate some of the mechanical load from reaching the nucleus. Indeed, the vimentin network (as labelled with vimentin-GFP overexpression) underwent larger deformations in cells with fibrillar adhesions (SI fig. 6). This protection also reduced the well-known effect of stretch-induced DNA damage^27–29^ as measured with the marker γH2AX (Fig. 5f,g and SI fig. 6).

To understand the mechanisms underpinning the protective effect afforded by the fibrillar adhesion – vimentin coupling, we imposed a 13% cell stretch in our computational model. In the condition where vimentin is anchored to the fibrillar adhesions, we observed high stresses in the extracellular matrix close to the adhesion sites (Fig. 5h). By assuming that this stress is frictional (due to the slip-bond nature of the adhesions as described in Methods) and thereby dissipates energy, this implies that vimentin anchoring to the fibrillar adhesions facilitates energy dissipation to the substrate. By contrast, vimentin decoupling from the fibrillar adhesions reduced stresses, energy dissipation, and therefore nuclear strain energy (from 2.5 to 1.9 fJ, Fig. 5h and methods). This in turn corresponds to a smaller nuclear deformation in cells with fibrillar adhesions (4%) than without (10%, Fig. 5c). Taken together, our experimental results in conjunction with a minimal component model reveal that vimentin anchoring to the ECM is crucial for effectively dissipating an applied stretching force and protecting the nucleus from deformation and damage.

## Discussion

Mechanotransduction processes include events that occur at the scale of seconds (such as integrin-mediated reinforcement^30,31^ or altered nucleocytoplasmic transport^3,8^), minutes (such as focal adhesion maturation^32^), hours (such as transcriptional responses or chromatin remodelling^28^) and days or longer (such as tumour growth^33,34^). Some of the longer-lived effects, such as chromatin reorganization or cell differentiation, can persist for some time after the mechanical stimulus has ceased, in a phenomenon that has been termed “mechanical memory”^35–37^. Even considering such memory effects, mechanical stimulation in physiological conditions may occur at all these timescales, potentially triggering constant, uncontrolled mechano-signalling. Just as biochemical signalling pathways do^38–41^, it is thus reasonable to hypothesize that mechanotransduction pathways should have mechanisms controlling the timescale of cellular response. To effectively control mechanotransduction, such mechanisms should in fact control not merely the timescale of cellular response, but of mechanical stimulation itself.

Here we demonstrate one such mechanism, by which matrix remodelling, fibrillar adhesion formation, and the anchoring of the vimentin network lock the nucleus in a mechanically deformed state. Effectively, this sets a low-pass filter to mechanical stimulation, setting the timescale of nuclear shape changes (and all subsequent mechanotransduction events) to the timescale of fibrillar adhesion remodelling (∼1h). This mechanism operates in response both to decreased and increased mechanical stimulation, and shows two major new properties of the ECM: i) it can regulate not only mechanosignalling but its timescale, and ii) this regulation depends not only on ECM protein composition^23^ and spatial organization^42^ as reported previously, but also on the conformation of its proteins (in this case, fibronectin). Our findings also reveal an unanticipated role of vimentin in maintaining rather than preventing nuclear deformation, as previously reported^25,26^.

Two relevant questions remain open. First, the molecular link between plectin 1f and the fibrillar adhesions remains unknown, and multiple molecular binding targets could potentially exist. Tensin is a possible candidate as it is a major component of the fibrillar adhesions^12^, and tensin-3 drives fibrillar adhesion formation via the interaction with talin^43^. Another candidate is Hic-5, which is fundamental for fibrillar adhesion formation and fibronectin remodelling via its interaction with tensin1^44,45^, and its ablation triggers vimentin network collapse^46^. Second, the fact that fibrillar adhesions are disrupted by latrunculin but not cytochalasin is intriguing. Potentially, this may be due to inaccessibility of cytochalasin to the barbed end of actin microfilaments within the long-lived fibrillar adhesion complexes (whereas latrunculin targets freely accessible actin monomers).

We anticipate that this mechanism could be relevant in physiological settings where slow and fast mechanical perturbations must elicit different cellular responses. For instance, fibroblasts in connective tissue associated with lungs, heart, the circulatory system, or the urinary system should respond to long-lasting mechanical alterations caused by wounding or tumour formation, but not to second-scale alterations caused by breathing, heart pumping, or bladder voiding. Whether such implications remain to be explored, the timescale regulatory mechanism unveiled here sets a foundation to explore such phenomena, with potential implications in homeostasis and disease.

## Acknowledgements

We thank S. Usieto, A. Menédez, N. Castro and M. Purciolas-Casas for providing technical support, A. Le Roux, L. Faure, A. Labernadie, R. Sunyer, J. Triadú-Gali, L. Rossetti and all members of the P. Roca-Cusachs and X. Trepat groups for invaluable discussions. A.E.M.B. is the recipient of a Sir Henry Wellcome fellowship (210887/Z/18/Z). P.R.-C. acknowledges funding from the Spanish Ministry of Science and Innovation (PID2019-110298GB-I00), the European Commission (H2020-FETPROACT-01-2016-731957), the Generalitat de Catalunya (2021 SGR 01425), The prize “ICREA Academia” for excellence in research, Fundació la Marató de TV3 (201936-30-31), and “la Caixa” Foundation (Agreement LCF/PR/HR20/52400004). IBEC is a recipient of a Severo Ochoa Award of Excellence from MINCIN. F.M.Y acknowledges funding from the National Institutes of Health (F31 DK126427). A.A-S is funded by the Spanish Ministry for Science and Innovation (FPU21/03952). K.S.A. acknowledges funding from the National Science Foundation (CBET 2033723) and the National Institutes of Health (R01 DK120921). The computational work was supported by National Cancer Institute awards U54CA261694 (V.B.S.); National Institute of Biomedical Imaging and Bioengineering awards R01EB017753 (V.B.S.) and R01EB030876 (V.B.S.); NSF Center for Engineering Mechanobiology GrantCMMI-154857 (V.B.S.); NSF Grant DMS-1953572 (V.B.S.). J.I. acknowledges funding from the Finnish Cancer Institute (K. Albin Johansson Professorship to J.I.) and Academy of Finland Centre of Excellence program (grant no. 346131)

## Author contributions

A.E.M.B and P.R.-C conceived the study. A.E.M.B and P.R.-C designed the experiments. A.E.M.B, A.A.-S, D.Z, and Z.K performed experiments. A.J. and V.B.S designed and implemented the computational model.

F.N.Y and K.B synthesised the photodegradable gel compounds and performed associated gel characterisation. I.A, I.G.-M, G.W X.T, J.I, K.S.A, V.B.S contributed to material, reagents, technical expertise and discussions. A.E.M.B and P.R.-C wrote the manuscript, with input from all other authors.

## Data availability

Data that support the findings of this study are in the article and supplementary data file. The other relevant raw data and source data are available from the corresponding author upon request.

## Code availability

Simulation codes are available upon request.

## Material and methods

### Cell Culture, transfections, and drug treatments

Human telomerase immortalized foreskin fibroblast (TIFFs) (a kind gift from J. Ivaska), were cultured in high glucose Dulbeccós modified eagle medium (DMEM, ThermoFisher Scientific) supplemented with 20% FBS (v/v, ThermoFisher), 2% Hepes 1M (v/v, H0887 Sigma) and 1% penicillin-streptomycin (v/v, 10378016 ThermoFisher). For stretch experiments, media was changed to CO_2_-independent media (18045054 ThermoFisher) supplemented with the same concentrations of FBS, Hepes, and penicillin-streptomycin as previous. Mammary epithelial cells (MCF10A) were purchased from ATCC, and cultured in DMEM-F12 (21331-020 LifeTechnologies) with 5% horse serum, 1% penicillin streptomycin, EGF (20 ng/ml), hydrocortisone (0.5 μg/ml), cholera toxin (100 ng/ml), and insulin (10 μg/ml). All cells were maintained at 37°^c^ with 5% CO_2_. Cell cultures were routinely checked for the presence of mycoplasma.

Transfections were conducted using the Neon transfection system (ThermoFisher Scientific) following the manufacturers instructions. TIFF cells were subjected to a single voltage pulse of 1650 V for a width of 20 ms. Cells were transfected the day before experiments, and cells were seeded ∼4hours prior to the experiment unless otherwise stated.

#### The plasmids used for transfections were

L_NLS-41-GFP mechano-reporter was generated in the lab from a previous study^8^. Plectin 1f-GFP and plectin 1f-8-GFP were generated from a previous study^47^. GFP was generated in the lab from previous study^3^. Membrane marker N-terminal Neuromodulin-GFP was a kind gift from Pr. F. Tebar. Tensin-1-eGFP was a kind gift from J. Ivaska. EGFP-Vimentin-7 was a gift from Michael Davidson (Addgene plasmid #56439; http://n2t.net/addgene:56439; RRID:Addgene_56439).

For the pharmaceutical inhibitor experiments, cells were seeded on fibronectin coated substrates for a minimum of 4 hours (unless otherwise stated) to allow fibrillar adhesion formation. All compounds were diluted and stored according to manufacturers instructions. Immediately prior to the experiments, compounds were diluted in cell culture media and warmed to 37°^c^ before adding to cells.

The drugs and concentrations used were: Blebbistatin (25 µM, B0560 Sigma), Cytochalasin D (1 µM, C2618 Sigma), Y-27632 (25 µM, 688001 Sigma), and LatrunculinA (0.5 µM, L5163 Sigma). Control cells were incubated with DMSO (Sigma), where the volume added was equal to the maximum volume of the drug conditions.

For the activation of integrin-α_5_β_1_ by Mn^2+^, after trypsin cells were resuspended in media containing 5mM Mn^2+^ and seeded onto 1.5kPa polyacrylamide gels. This concentration of Mn^2+^ was maintained throughout the duration of the experiment.

### Fibril blocking approaches

The PUR4 (also known as FUD) peptide (sequence: KDQSPLAGESGETEYITEVYGNQQNPVDIDKKLPNETGFSGNMVETEDT) and the scrambled control (sequence: EKGYSKPPVGNEGGDQVDEYDTMSQTKLEDEGNTLISPITFENATEQVN) were synthesised by ThermoFisher Scientific without any tags or modifications. In all experiments, after trypsin the cells were resuspended in media containing the PUR4 or scramble peptide at a final concentration of 500nM. The peptide was maintained in the media for the duration of the experiment.

For blocking of the integrin α_5_β_1_, cells in suspension were incubated for 20 minutes at 37°^c^ in the blocking antibody (integrin α_5_β_1_, clone JBS5, Sigma) or control antibody (IgG) before seeding onto 10µg/mL fibronectin coated glass surfaces. Both blocking antibodies were used at concentration of 10µg/mL.

Gluteraldehyde blocked surfaces were prepared as described previously^19^. Briefly, glass surfaces were coated with 10μg/ml fibronectin overnight at 4°C or 1 hour at 37°C. Surfaces were then treated with a 1% gluteraldehyde (Sigma Aldrich) in milliQ H_2_O solution for 10 minutes at room temperature. Surfaces were then thoroughly rinsed with H_2_O and left to incubate in freshly prepared 1% BSA (Sigma Aldrich) solution for at least 20 minutes at 37°C before cell seeding.

### Immunostainings

Cells were fixed with 4% paraformaldehyde for 10 minutes at room temperature and rinsed thrice with PBS. Cells were permeabilized with 0.1% Triton-X for 5 minutes and then blocked with 0.5% Fish-Gelatin (SigmaAldrich) for 1 hour (except manganese treated cells which were permeabilized using 0.5% TritonX for 15 minutes). Cells were incubated with the primary antibody for 1 hour diluted in the 0.5% fish gelatin blocking solution, washed with blocking solution for 30 minutes, and incubated with the secondary antibody labelled with Alexa fluorophore (ThermoFisher, 1:300 dilution) for 1 hour. In the case of actin staining, phalloidin (SigmaAldrich, 1:1000) was added with the secondary antibody. Hoechst (1:2000) was added for 5 minutes to label the nuclei, and samples were washed thoroughly.

#### The primary antibodies and their dilutions were

YAP (1:300, sc-101199, Santa Cruz) or (1:300, 14074S, Cell Signaling). Integrin α5, clone Snaka51 (1:300, MABT201, Millipore). LaminB (1:300 ab16048, abcam). Paxillin (1:300, ab32084, abcam). Twist (1:100, SC-81417, Santa Cruz). Snail (1:50, Ab224731, abcam). Tensin-1 (1:200, ab233133, abcam). Fibronectin (1:300, F3648, Sigma). Vimentin (1:600, ab92547, abcam). γH2Ax (1:300, 2577, Cell Signaling).

### Polyacrylamide Gel

Polyacrylamide gels of variable rigidity were prepared as previously described^8^. Briefly, glass bottom dishes (MatTek) or glass coverslides were treated a solution of 3-(Trimethoxysilyl)propyl methacrylate (Sigma), acetic acid, and 96% ethanol (1:1:14) for a minimum of 10 minutes. The glass was then thoroughly rinsed in 96% ethanol and dried. Gels were prepared by mixing different concentrations of acrylamide and bis-acrylamide to produce gels of different rigidity according to previous characterisation^8^, with 2% v/v 200-nm-diameter fluorescent carboxylate-modified beads (Fluospheres, Thermo Fisher Scientific), 0.05% v/v ammonium persulfate (Sigma Aldrich) and 0.05% tetramethylethylenediamine (Sigma Aldrich), in PBS 1×. To cast the gels, 22 μl was placed on top of the treated glass and then covered with an 18 mm circular coverslip. Gels were left for 45 min to polymerize at room temperature. Finally, gels were submerged in PBS 1× and the top coverslip was removed. To coat gels, we prepared a mixture containing 10% HEPES pH (0.5M pH6.0), 0.004% Bis-acrylamide, 0.05% Igracure 2959 and 4% Acrylic-acid NHS/DMSO (10 mg/ml, A8060 Sigma) in milliQ water. Gels were coated in this mixture and then illuminated with UV light for 8 minutes. Gels were then washed twice in 50mM HEPES pH7 and twice in PBS 1X, and incubated with 10ug/mL fibronectin in PBS overnight at 4°C, sterilised by UV-treatment in a laminar flow hood, washed once with PBS and immediately used.

### Photodegradeable compound synthesis

Photodegradable precursors were prepared as previously described^22^. Briefly, the acrylate functionalized photodegradable monomer was synthesized by suspending 4-[4-(1-hydroxyethyl)-2-methoxy-5-nitrophenoxy]butyric acid (0.0166 mol, Sigma-Aldrich) in anhydrous DCM (90 mL). The mixture was purged with argon; triethylamine (0.0664 mol) was added to the flask by syringe; and acryloyl chloride (0.0547 mol) in dry DCM was added dropwise at 0°C. The reaction was kept under argon atmosphere and allowed to proceed overnight at room temperature. The reaction mixture was then added to DI water (0.5 L) and allowed to stir for 2 hours at room temperature, before being extracted with chloroform (5 x 200 mL washes). The organic phase was dried over NaSO_4_ and concentrated by rotary evaporation to obtain the acrylate functionalized photodegradable crosslinker.

To synthesize the photodegradable PEG crosslinker (PEGdiPDA), the acrylate functionalized photodegradable monomer (6 mmol) was dissolved in NMP (15 mL) and purged with argon. The coupling agent 2-(1H-benzotriazole-1-yl)-1,1,3,3-tetramethyluronium hexafluorophosphate (HBTU, 6.6 mmol), 1-hydroxybenzotriazole (HOBt, 6.6 mmol), and diisopropylethylamine (DIEA, 0.012 mol) were then added to the reaction mixture and stirred for 5 minutes before the addition of PEGdiamine (0.6 mmol, 2 kDa) in NMP. The reaction mixture was heated and vortexed until all reactants had completely dissolved, and left to stir overnight at room temperature. The reaction mixture was then precipitated in diethyl ether at 0°C and collected by centrifugation. The macromer product was redissolved in water and centrifuged (21,000 RPM, 1 hr) to yield a dark brown pellet with a clear supernatant. The supernatant was collected, dialyzed (SpectraPor 7, CO 1000 g/mol), and lyophilized to produce a white powder (39 % yield) that was used in experiments.

### Characterization of photodegradable gel mechanical properties

Photodegradable gels were prepared by first mixing 5.4wt% PEGdiPDA, 9.6wt% PEG400acrylate, and 6.6mM sodium acrylate in PBS before degassing for 5 minutes on ice. Polymerization was initiated by addition of 200mM TEMED and 100mM APS, which were pre-incubated on ice, and drops were added between Sigmacote (Sigma-Aldrich) treated glass slides with either 200 µm spacers for 12 µL gels or 100 µm spacers for 6 µL gels. The gels were left to polymerize for 10 minutes before the top glass slide was removed and the hydrogels were transferred to a well plate with PBS (500 µL). Following equilibration for 30 minutes in PBS, the hydrogels were transferred to a rheometer (DHR-3, TA Instruments) equipped with a light curing accessory (Omnicure 1000, Lumen Dynamics) and an 8 mm parallel plate tool. The 6 µL gels were used to track in situ network evolution during irradiation (365 nm, 10mW cm^-2^), and the 12 µL gels were used to evaluate rheological properties of equilibrium swollen samples before and after pre-selected doses of irradiation. All rheological characterization experiments utilized a strain of 1% and a frequency of 1 Hz.

### Photodegradable gel

Glass bottom dishes were activated using the same protocol as the polyacrylamide gels. Photodegradable gels were prepared by first mixing 5.4wt% PEGdiPDA, 9.6wt% PEG400acrylate, and 6.6mM sodium acrylate in PBS before degassing for 5 minutes on ice. Polymerization was triggered by the addition of 5% TEMED and 10% APS (2M), which were pre-incubated on ice, and a 22μL drop of gel mixture was placed in the centre of the glass bottom dish and covered with a 18mm coverslip to achieve uniform spreading. The gels were left to polymerise for 10 minutes before the addition of PBS and the removal of the top coverslip.

For functionalisation, we prepared a mixture containing 100mM 1-Ethyl-3-(3’-dimethylaminopropyl)carbodiimide HCL (8510070025, Sigma), and 200mM N-Hydroxysuccinimide (130672, Sigma) in 20mM HEPES buffer pH7. Gels were incubated in this mixture for 20 minutes at 37°C. The gels were rinsed once with HEPES buffer and once with PBS. The gels were then incubated with 10µg/mL of fibronectin overnight at 4°C. To initiate gel softening, gels were placed under a UV lamp (UVP 365nm, 15 Watt) for 4.5 minutes.

### Image acquisition

Epifluorescent images and time lapse microscopy was performed with inverted microscopes (Nikon Eclipse Ti) equipped with thermal, CO_2_ and humidity control. Microscopes were equipped with an ORCA Flash4.0 camera (Hamamatsu) and controlled with MetaMorph (version 7.7.1.0) or Micromanager. Most images were taken with a 60X objective (plan apo; NA 1.2; water immersion) unless otherwise stated.

For time-lapse acquisition of the change in mechano-reporter L_NLS-41-GFP localisation with drug treatments, a single image was acquired prior to pharmacological treatment, and then images were acquired every 5 minutes for a total duration of 1 hour. For time-lapse acquisition of the change in mechano-reporter L_NLS-41-GFP localisation upon *in-situ* photodegradable gel softening, images were taken on a Nikon TiE inverted microscope equipped with a spinning disk confocal unit (Andor) and a Sona sCMOS camera (Andor), using a 40X objective (Plan Fluor; NA 0.75) controlled with Fusion software. A single z-stack was acquired prior to UV softening. Gels were then softened for 4.5minutes with a UV lamp, and z-stack images were taken every 10 minutes for 2 hours.

Confocal images of nuclear height and plectin1F were acquired a Zeiss LSM880 inverted confocal microscope using Zeiss ZEN2.3 SP1 FP3 (black, version 14.0.24.201), using a 63X 1.46 NA oil immersion objective. Confocal images of the vimentin network were taken using a Nikon TiE inverted microscope with a spinning disk confocal unit (CSU-WD, Yokogawa) and a Zyla 4.2 sCMOS camera (Andor) using a 60X objective (plan apo; NA 1.2; water immersion) controlled with Micromanager.

### Traction force microscopy

Traction force microscopy experiments were performed as described previously^48^. Briefly, cells were seeded on 15kPa polyacrylamide gels embedded with fluorescent beads. Images of the cells and the beads were acquired prior to pharmacological treatment, and 30 minutes after pharmacological treatment. Cells were then removed from the gel using Trypsin to obtain a reference image of the beads. Local gel deformation was computed with a custom particle imaging velocimetry (PIV) software^49^ in Matlab (MathWorks Inc.). Traction forces were computed using Fourier traction microscopy with a finite gel thickness and the mean of each cell was calculated.

### Cell stretch experiments and quantification

Stretchable polydimethylsiloxane (PDMS) (Sylgard 184 Silicone Elastomer Kit, Dow Corning) membranes were prepared as previously described^30^. Briefly, a mix of 10:1 base to crosslinker was spun for 1 min at 500 rpm and cured at 65 °C overnight on plastic supports. Once polymerized, membranes were peeled off and assembled onto the stretching device. The PDMS membranes were functionalized with 10 µg/mL fibronectin overnight at 4 °C. TIFF cells were seeded for at least 4 hours (unless otherwise stated) prior to the stretch experiment. Immediately before stretch, the cell media was changed to CO_2_-independent media. The stretch experiments were performed by mounting the stretching device on an upright microscope (Nikon eclipse Ni-U) equipped with temperature control and controlled with Metamorph. Calibration of the system was performed using PDMS coated with fluorescent beads, to ensure that the vacuum applied a 10% stretch to the PDMS membrane. Each membrane was stretched for a maximum of 6 times per experiment. The percentage change in area of the nucleus and cell membrane upon stretch was calculated by segmenting the fluorescent signal from the Hoechst or membrane marker respectively, before and during stretch. For DNA damage experiments, cells were subjected to 5 cycles of 30 seconds of 10% stretch, 10 seconds of release. The stretch system was immediately removed from the microscope and the cells were fixed and stained with γH2Ax and Hoechst. The Hoechst signal was used to segment the nuclei, and the mean intensity of each nucleus was measured correcting for the background.

### Adhesion disassembly times

To measure the disassembly times of focal adhesions compared to fibrillar adhesions, TIFF cells were transfected with either paxillin-GFP or tensin1-GFP, respectively. Transfections were performed 24 hours prior to the experiment. On the day of the experiment, cells were seeded on fibronectin-coated glass bottom dishes (Mattek) and left to form adhesions for a minimum of 4 hours. Adhesion dynamics were acquired using a Zeiss LSM880 inverted confocal microscope with a 63X 1.46NA oil immersion objective. For cells expressing paxillin-GFP, images were acquired every 120 seconds for approximately 2.5 hours. For cells expressing tensin1-GFP, images were acquired every 300 seconds for approximately 10 hours. The intensity of an adhesion was tracked with time, from the initial formation until disappearance. The plot of adhesion intensity was then fit with a gaussian, and the disassembly time was measured at the time from the gaussian peak until the return to background levels.

### Image analysis

#### Nuclear / Cytoplasmic (N/C) ratio analysis

The nuclear/cytoplasmic ratio was quantified by measuring the mean fluorescence intensity of a nuclear region (I_nucleus_) and the intensity of a cytoplasmic region immediately adjacent (I_cytoplasm_). The nuclear region was determined from the Hoechst staining. The ratio was calculated using the following formula:

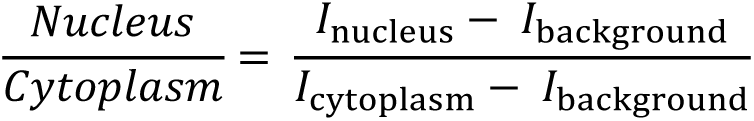

Where I_background_ is the mean fluorescence intensity of a region outside of the cell. For the quantification of the mechano-reporter L_NLS-41-GFP with time, the N/C ratio was calculated at each time point. For all drug treatment experiments, the N/C ratio at each timepoint was normalized to the N/C ratio prior to the addition of the compound. For quantification of the N/C mechano-reporter L_NLS-41-GFP ratio during *in-situ* gel softening experiments, a single confocal plane was selected and the N/C ratio was normalized between the pre-softened N/C ratio and the ratio at the final timepoint.

#### Fibrillar adhesion quantification

Fibrillar adhesions were marked with the Integrin α5, clone Snaka51 antibody (or tensin-1 antibody for the blocking antibody experiments). To quantify the extent of fibrillar adhesion formation, the fibrils in the area under the nucleus (determined from Hoechst staining) were detected using the Fiji Ridge Detection plugin, and the percentage area occupied by fibrils was computed. For cells seeded on soft gels, instead the length of the fibrillar adhesions was calculated to circumvent changes in focal plane across the whole cell. For each cell, the length of ∼5 representative fibrillar adhesions under the nucleus were measured.

#### Focal adhesion length

Focal adhesion length was obtained by measuring the length of ∼5 representative focal adhesions at the cell periphery for each cell.

#### Nuclear height

Nuclear height was measured from z-stack confocal images of laminB stained nuclei. Each nucleus was resliced along the long-axis, an intensity profile was created, and the height was measured from the distance between the two peaks of maximum laminB intensity.

#### Vimentin spreading

To calculate the area occupied by vimentin, confocal stacks were acquired for cells stained with actin and vimentin. The area of the actin and vimentin network was calculated by thresholding the z-projection (sum) of each channel. The percentage area of the vimentin network with respect to the total cell area (from the actin network) was computed for each cell.

#### Actin Anisotropy

The actin anisotropy was analysed using the FibrilTool plugin in imageJ^50^.

### Computational model

#### Constitutive Models for the cell, nucleus and substrate

To fully describe the effect of mechanical stress (generated due to cellular contraction and/or applied stretch) on the nucleus, we consider the following key components in our computational model: i) contraction due to myosin motors (red, Figure 3n), ii) actin filaments blue, Figure 3n), iii) vimentin intermediate filaments (green, Figure 3n), iv) microtubules and v) fibrillar adhesions. In our model, the cell cytoskeleton is considered to consist of spatially varying representative volume elements (RVEs), each of which is comprised of the components (i-iv) described above (Figure S7(a)). We assume initially uniform and isotropic distribution of these elements and describe how, due to the action of contractile forces and the resulting stress field, these cytoskeletal components are redistributed in a more anisotropic manner, facilitating force transfer from the cell cytoskeleton to the nucleus. Also, the extracellular matrix is modelled as a linear elastic material with elastic modulus 70kPa, while the nucleus is similarly modelled as an elastic material with Young’s modulus and shear modulus values as listed in Table 1. We describe each of these components here.

**Table 1.** List of parameters used in the simulations.

| Parameter | Description | Value |
| --- | --- | --- |
| $K_{nuc}$ | Bulk modulus of nucleus | 2.5 kPa |
| $\mu_{nuc}$ | Shear modulus of nucleus | 1.2 kPa |
| $E_{cell}$ | Elastic modulus of cell at 0.1% strain | 2.1 kPa |
| $C_1$ | Mooney-Rivlin Parameter | 0.28 Pa |
| $C_2$ | Mooney-Rivlin Parameter | 700 Pa |
| $\eta_d$ | Frictional viscosity constant | 90 nN.s/$\mu\text{m}$ |

*Cytoskeletal contraction due to myosin molecular motors.* Myosin motors are treated as force dipoles (pair of equal, but oppositely oriented forces) that bind to actin filaments and generate cellular contractility^51^ (Figure 3n, inset in green). The volume-averaged density of bound motors can be represented by a symmetric tensor, 𝜌_𝑖𝑗_, whose components represent cytoskeletal contractility along different directions^52^. Within our coarse-grained approach, the contraction due to myosin motors is represented as an isotropic stress tensor (ρ_11_=ρ_22_=ρ_33_=ρ) of magnitude 1.5kPa, applied at every point in the cell cytoskeleton. Due to cytoskeletal contraction, compressive stress 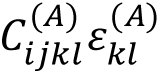 are generated in components in compression (like vimentin), while tensile stresses *σ*_𝑖𝑗_ are generated in the cytoskeletal components under tensile strain (actin filaments). By force balance, the contractility is given as:

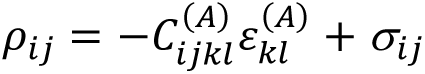

where 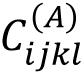 and 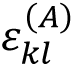 are the stiffness tensor and strain of the components in compression (SI Fig. 7), namely the microtubules and vimentin.

*Actin Filaments and actin-vimentin interactions.* The actin filaments experience tension as the cell contracts and hence are in series with the myosin element. Vimentin intermediate filaments (VIF) interact with actin through direct physical contact facilitated by cross-linkers and direct binding^53^. Hence, the vimentin intermediate filaments in contact with actin also experience tensile stresses and is added in series to the non-linear elastic element representing actin filaments (SI Fig. 7).

*Vimentin-microtubule network under compression.* Vimentin intermediate filaments near the perinuclear region interact with microtubule elements and are experimentally reported to stabilize them^54,55^. To represent this effect, we add another set of vimentin elements in parallel with the microtubules in compression. Hence, there are two sets of vimentin intermediate filaments, one that is in direct physical contact with actin and under tension, while the other is enmeshed with microtubules under compression, reinforcing them^56^. Also, vimentin intermediate filaments have been observed to stiffen under compressive strains, leading to an overall compressive stiffening of cytoskeletal networks^26^. To represent the above effects, the cytoskeleton is modelled as a nearly incompressible, hyperelastic solid that stiffens under compressive strains. First, we define the deformation gradient 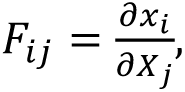 as a second order tensor that maps infinitesimal line elements 𝑑𝑿 in the reference configuration to corresponding infinitesimal line elements 𝑑𝒙 in the current configuration. Further, we define 𝑪 = 𝑭^𝑻^𝑭 to be the right Cauchy Green deformation tensor whose normal components represent stretch along a given direction, while shear components represent change of angle. A Mooney-Rivlin constitutive equation is used to represent this stiffening behaviour and the strain energy of the cell can be defined as:

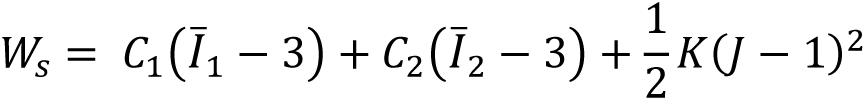

In the above equation, the Jacobian 𝐽 = det(𝑭) the determinant of the deformation gradient tensor, while 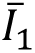 and 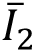 are the first and second invariants of the deviatoric part of 𝑪, respectively. The parameters C_1_ and C_2_ are Mooney-Rivlin parameters while 𝐾 is the bulk modulus of the cell. In the limit of small strains, the parameters 𝐶_1_ and 𝐶_2_can be related to the shear modulus of the cell µ as : µ = 2(𝐶_1_ + 𝐶_2_). The values of these parameters are listed in Table 1 and the elastic modulus of the cell for compressive strains in the range of 0.001 to 0.5 are found to be 2.1-2.9 kPa, which provides reasonable agreement with the Young’s modulus measured for living fibroblasts using AFM^57^. The high levels of compressive strain near the nucleus due to the contractile stress leads to the formation of a stiff region representing the vimentin cage observed experimentally (see principal stress *σ*_3_, SI Figure 7). By contrast, in the cell cytoskeleton, high tensile stresses are observed close to the basal plane, particularly near the cell periphery (see principal stress *σ*_1_, SI Figure 7) representing the actin and vimentin intermediate filaments in tension.

#### Adhesive and frictional forces due to fibrillar adhesions

Due to cytoskeletal contraction, the vimentin cage around the nucleus is gradually pushed down and is anchored by fibrillar adhesions that are present near the centre of the cell. We propose that the fibrillar adhesions exert an adhesive force fa on the edge of the vimentin cage (Figure S3 (A)) once they are engaged, which prevents nuclear rebound upon loss of contractile forces. We model this by applying a vertical force along the vimentin edges which remain even when contractile forces are removed. We estimate this force to be of the same magnitude as the contractile force needed to push down the vimentin cage.

Fibrillar adhesions are defined by α5β1-integrin and tensin family of proteins which form bonds with ligands on the ECM. On the application of stretch, the bond between the fibrillar adhesions and the ligands on the ECM (a polymer structure) are ruptured. This rupture corresponds to overcoming of energy barriers by thermal activation (note that, we assume that the vimentin remains anchored to the fibrillar adhesions as it slides). The disengagement of the fibrillar adhesion-ligand bond follows a Hill Type relation, where the velocity of the sliding adhesions decays exponentially as the activation energy associated with bond rupture 𝐸_𝐴_ increases:

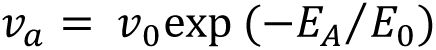

where 𝑣_0_ is the maximum sliding velocity (when the activation energy 𝐸_𝐴_ is zero) and 𝐸_0_is an energy scale related to thermal or active noise. The activation energy can be expressed as the work done by a dissipative force 𝑓_𝑑_ in translocating the fibrillar adhesions by a molecular sliding distance 𝑎:

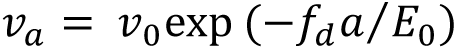

Assuming that the activation energy associated with the rupture is much smaller than 𝐸_0_, the above equation can be linearized, and expressed as:

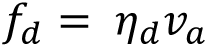

where, 𝜂_𝑑_ = 𝐸_0_/𝑎𝑣_0_ is a frictional dissipative constant that is inversely related to the mobility of individual fibrillar adhesions. Setting η_d_ as zero is equivalent to the case of fibrillar adhesions not anchored to the ECM and is representative of cells lacking fibrillar adhesions.

#### Geometry, Mesh and Boundary Conditions

The model for the cell cytoskeleton, nucleus and the fibrillar adhesions is implemented in COMSOL Multiphysics, within a finite element framework. The cell is modelled as an ellipsoid of semi-axes lengths 15 and 12µm, while the nucleus is modelled as a spheroid of radius 3.7 µm located at the centre of the cell. The substrate is modelled as a rigid cylinder of radius 50 µm. Due to rotational symmetry of the cell-substrate system, an axisymmetric analysis is conducted, with horizontal roller boundary conditions applied to the top and bottom ends of the substrate. While the cell and nucleus fully rest on the substrate, a small hemispherical region around the nucleus 0.5 µm thick and separated from the nucleus by 0.1 microns is initially separated by a gap of 0.03 µm from the substrate. This represents the vimentin cage that forms upon the action of contractile forces and is pushed down and eventually is anchored to the fibrillar adhesions. Contact conditions are implemented at the vimentin-nucleus interface representing the effect of nesprins and other molecules that directly transfer contractile stresses to the nucleus. In addition, contact conditions are implemented at the vimentin-substrate interface to represent the adhesion between the vimentin and the fibrillar adhesions near the centre of the cell. Triangular mesh elements are used to discretize the cell geometry with a minimum element size of 0.001 µm near the contact zones to accurately resolve the stresses and displacements at contact.

### Material and software availability

This study did not generate new reagents.

### Statistics

Statistical analyses were performed with GraphPad Prism software (GraphPad, version 9). Statistical significance was determined using the tests indicated in each figure legend. The number of data points and number of independent experiments performed for each experiment is stated in the figure legend.

