## Supplemental file for "Fibrillar adhesion dynamics govern the timescales of nuclear mechano-response via the vimentin cytoskeleton"

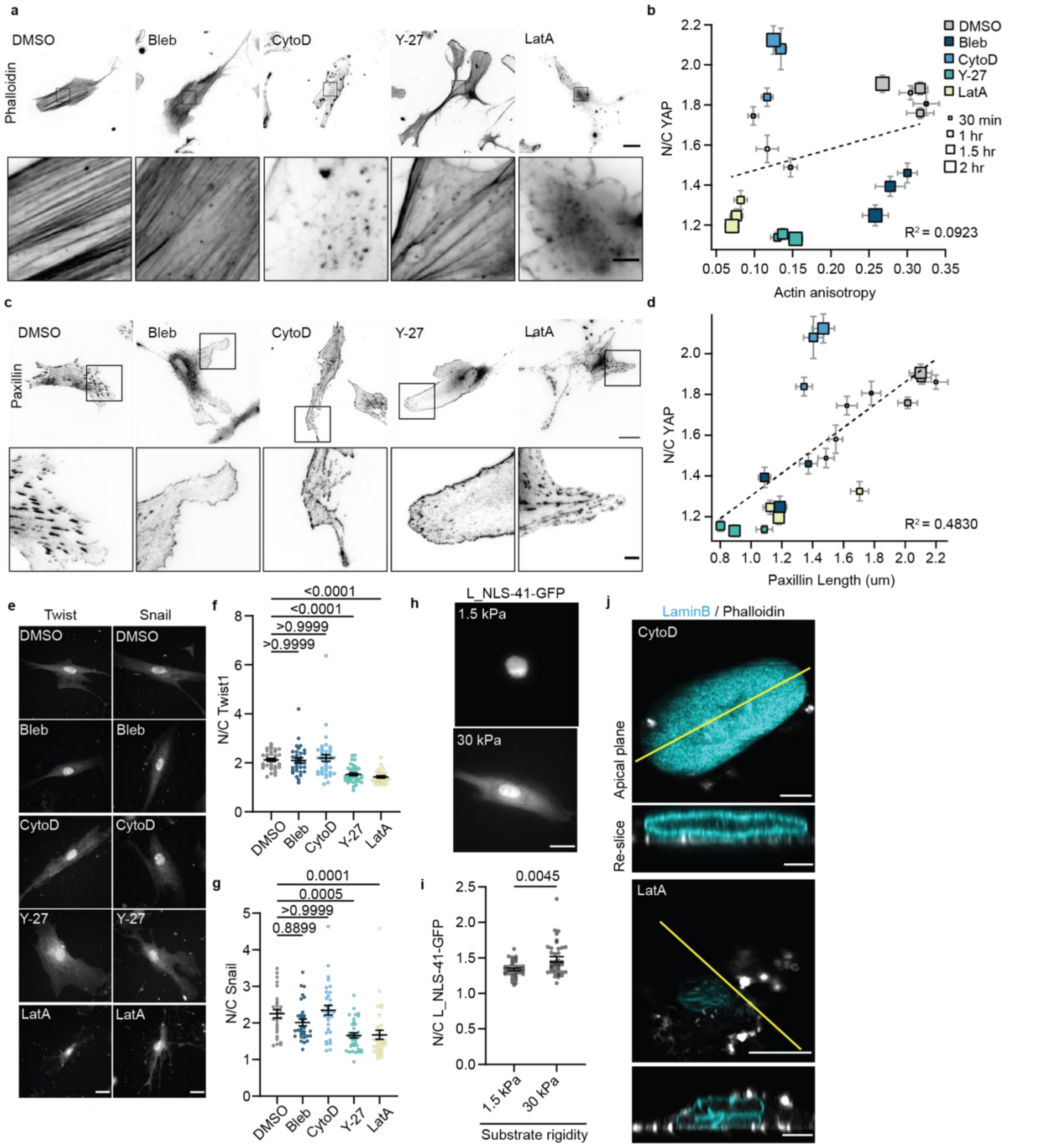


**Supplementary figure 1. Additional characterization of the effect of pharmacological treatment. a.** Images of the actin cytoskeleton stained with phalloidin, after 2 hour treatment with different contractility inhibiting drugs; scale bar 25 µm (5 µm in zoomed inset). **b.** Correlation between the N/C YAP ratio and the actin anisotropy for different drug treatments (colour coded) with different incubation time (size coded). 3 independent experiments with a minimum of 41 cells per condition. **c.** Images of focal adhesions (stained with paxillin) after 2 hour treatment with different pharmacological treatment; scale bar 25 µm (5 µm in zoomed inset). **d.** Correlation between the N/C YAP ratio and paxillin length for the different drugs at different incubation times. **e.** Immunostainings of Twist and Snail localization after 30 minute pharmacological treatment. Scale bar 25µm. **f.** Quantification of N/C twist ratio upon a 30 minute treatment with different pharmacological treatments. (n=33/32/38/36/32 cells for DMSO/bleb/cytoD/Y-27/latA from 2 independent experiments; Kruskal-Wallis test with Dunn's multiple comparison test). **g.** Quantification of N/C snail ratio upon a 30 minute treatment with different pharmacological treatments. (n=28/32/32/35/32 cells for DMSO/bleb/cytoD/Y-27/latA from 2 independent experiments; Kruskal-Wallis test with Dunn's multiple comparison test). **h.** Images of L_NLS-41-GFP transfected cells seeded on 1.5 kPa or 30 kPa gels. Scale bar 25µm. **i.** Quantification of N/C L_NLS-41-GFP expressing cells on 1.5 kPa and 30 kPa gels. (1.5 kPa, n=36 cells; 30 kPa, n=40 cells from 2 independent experiments; Mann-Whitney test). **j.** Confocal fluorescence images of the nucleus labelled with laminB (cyan) and the actin cytoskeleton labelled with phalloidin (grey) after 1 hour pharmacological treatment with CytoD or LatA. The XY image is a single slice at the apical plane on top of the nucleus (scale bar is 10 µm). The re-slice is taken at the yellow line (scale bar is 5 µm).

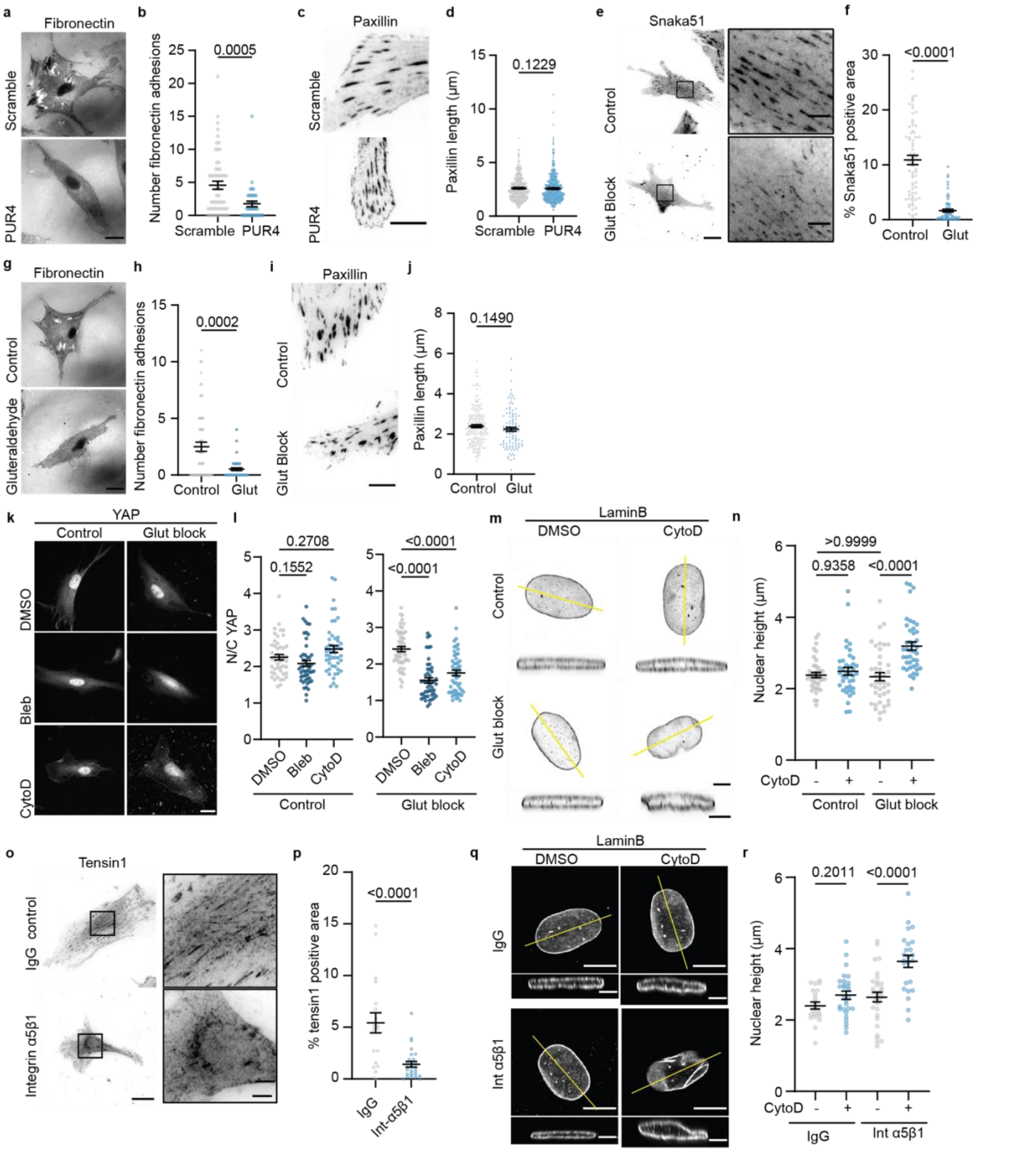


**Supplementary figure 2. Additional characterization of the inhibition of fibrillar adhesion formation and the impact for cellular response upon contractility inhibition. a.** Images of fibronectin stainings for cells seeded in scrambled or PUR4 peptide. Scale bar 25 µm. **b.** Quantification of number of fibronectin adhesions for scramble or PUR4 conditions. Adhesions were only considered if the intensity was above the background fibronectin intensity. P=0.0005 calculated with Mann-Whitney test scramble, n=58 cells and PUR4 n=44 cells from 2 independent experiments. **c.** Images of paxillin stained focal adhesions in cells with scrambled or PUR4 peptide. Scale bar 10 µm. **d.** Analysis of paxillin adhesion length for cells seeded with the scrambled peptide or the PUR4 peptide. P=0.1229 calculated with Mann-Whitney test, scramble n=440 adhesions, PUR4 n=524 adhesions from 4 independent experiments. **e.** Images of cells seeded on control and gluteraldehyde blocked glass substrates stained with the integrin α5 antibody, clone Snaka51. Scale bar 25 µm (zoom insert, 5 µm). **f.** Analysis of the percentage area under the nucleus that is occupied by fibrillar adhesions for cells seeded on control or gluteraldehyde blocked glass substrates. (p<0.0001 calculated with Mann-Whitney test, control n=69 cells and glut n=68 cells from 5 independent experiments). **g.** Images of fibronectin stainings for cells seeded in control or gluteraldehyde block conditions. Scale bar 25 µm. **h.** Analysis of number of fibronectin adhesions for the control and glut blocked condition. Adhesions were only considered if the intensity was above the background fibronectin intensity. (p=0.0002 calculated with Mann-Whitney test, control n=61 and glut n=62 cells from 2 independent experiments). **i.** Images of paxillin stained cells seeded on control and gluteraldehyde blocked glass substrates. Scale bar 10 µm. **j.** Quantification of paxillin adhesion length in cells seeded on control and gluteraldehyde blocked glass substrates. (p=0.1490 calculated with Mann-Whitney test, control n=175 adhesions and glut n=105 adhesions from 2 independent experiments). **k.** Example YAP stained cells after 30 min pharmacological treatment with control or gluteraldehyde blocked substrates. Scale bar 20 µm. **l.** Quantification of N/C YAP ratio for control cells or cells seeded on gluteraldehyde blocked substrates and subjected to 30min pharmacological treatment. (Control; n=47/49/49 cells and Glut; n=52/48/53 cells for DMSO/bleb/cytoD from 3 independent experiments; Kruskal-Wallis test with Dunn's multiple comparison test). **m.** Example images of LaminB stained nuclei for cells on control or glut blocked substrates treated with DMSO or CytoD. Scale bar 5 µm. **n.** Quantification of nuclear height of cells on control or glut blocked substrates treated with DMSO or cytoD. Control. (control; DMSO n=40, cytoD n=39 cells. Glut; DMSO n=42, cytoD n=42 cells from minimum of 3 independent experiments; 2-way ANOVA with Tukey's multiple comparison test). **o.** Immunofluorescent images of cells marked with tensin1 in cells treated with IgG control antibody or Integrin α5β1 (clone JBS5) blocking antibody (scale bar 25 µm). Black box indicates zoomed image, scale bar 10 µm. Fibrillar adhesions were marked with tensin1 rather than integrin α5 antibody (clone Snaka51) due to antibody compatibility conflicts as both primary antibodies are raised in mouse. **p.** Analysis of the percentage area of tensin1 under the nucleus for cells treated with a control blocking antibody or integrin α5β1 blocking antibody. P<0.0001 calculated with Mann-Whitney test from control n=19 cells and integrin α5β1 n=27 cells, from 2 independent experiments. **q.** Example fluorescence images of nuclei labelled with laminB in control IgG antibody blocked cells or integrin α5β1 blocked cells treated with DMSO or cytoD. Central plane of the nucleus, scale bar 10µm. Reslice of the nuclei corresponding to the yellow time position, scale bar 5µm. **r.** Analysis of nuclear height for IgG blocked (grey) or integrin α5β1 blocked (blue) cells treated with DMSO or cytoD. P-values calculated with two-way ANOVA with Tukey correction for multiple comparisons from 3 independent experiments (IgG DMSO, n=22 cells; IgG cytoD, n=28; α5β1 DMSO, n=32; α5β1 cytoD, n=24).

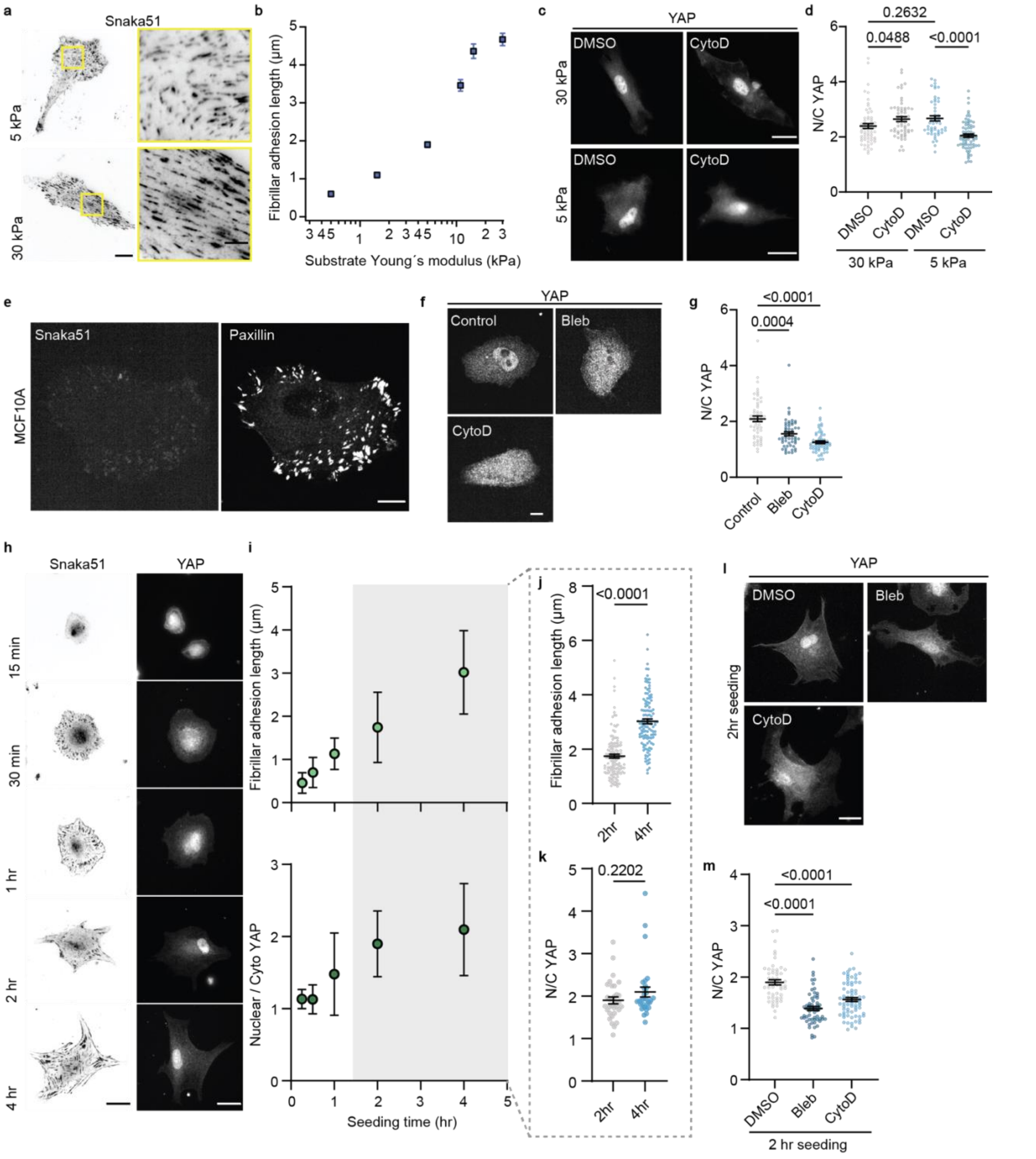


**Supplementary figure 3. Additional approaches to inhibiting the formation of fibrillar adhesions. a.** Example immunofluorescence images of cells marked with snaka51 antibody seeded on 5kPa and 30kPa polyacrylamide gels for a minimum for 4 hours. Scale bar 25 µm / zoom 5 µm. **b.** Quantification of fibrillar adhesion length as a function of substrate rigidity. Each data point contains a minimum of 64 fibrillar adhesions from 2 independent experiments. **c.** Immunostaining images of YAP in cells seeded on 30 kPa or 5 kPa gels, treated with DMSO or cytoD.

Scale bars 25 µm. **d.** Analysis of N/C YAP ratio in cells seeded on 30 kPa or 5 kPa gels and treated with DMSO or cytoD (30 kPa DMSO, n=61; 30 kPa cytoD, n=56; 5 kPa DMSO, n=52; 5 kPa cytoD, n=73; two-way ANOVA with Tukey multiple comparison test from 3 independent experiments). **e.** Example images of MCF10A cells stained with Snaka51 for fibrillar adhesions and Paxillin reveal the absence of fibrillar adhesions. Scale bar 15µm. **f.** Example YAP images of MCF10A cells after 30 minutes in the indicated pharmacological treatment. Scale bar 15µm. **g.** Quantification of N/C YAP ratios for control cells, blebbistatin, or cytochalasinD treated MCF10A cells. (n=54/53/60 cells for DMSO/bleb/cytoD from 2 independent experiments; Kruskal-Wallis test with Dunn's multiple comparison test). **h.** Example immunofluorescent images of cells seeded after indicated seeding time. Cells marked with integrin α5, clone snaka51 and YAP. Scale bar 25 µm. **i.** Analysis of the length of the fibrillar adhesions as a function of seeding time on fibronectin coated glass substrates (upper panel). Data from 2 independent experiments (15min, n=89 adhesions; 30min, n=115; 1hr, n=124; 2hr, n=130; 4hr, n=120). Analysis of the N/C YAP ratio as a function of seeding time (lower panel). Data from 2 independent experiments (15min, n=30 cells; 30min, n=29; 1hr, n=33; 2hr, n=32; 4hr n=31). Error bars are standard deviation. **j.** Quantification of fibrillar adhesion length corresponding to the two seeding times 2hr and 4hr. (n=130/120 adhesions for 2hr/4hr from 2 independent experiments; Mann-Whitney test). **k.** Analysis of N/C YAP ratio at 2hr and 4hr seeding time. (n=23/31 cells for 2hr/4hr from 2 independent experiments; Mann-Whitney test). **l.** Example images of YAP stained cells seeded for 2 hours and treated with DMSO, blebbistatin or cytochalasinD. Scale bar is 25 µm. **m.** Quantification of N/C YAP ratio in cells seeded for 2 hours before a 30 minute drug treatment. (n=54/61/73 cells for DMSO/bleb/cytoD from 3 independent experiments; Kruskal-Wallis test with Dunn's multiple comparison test).

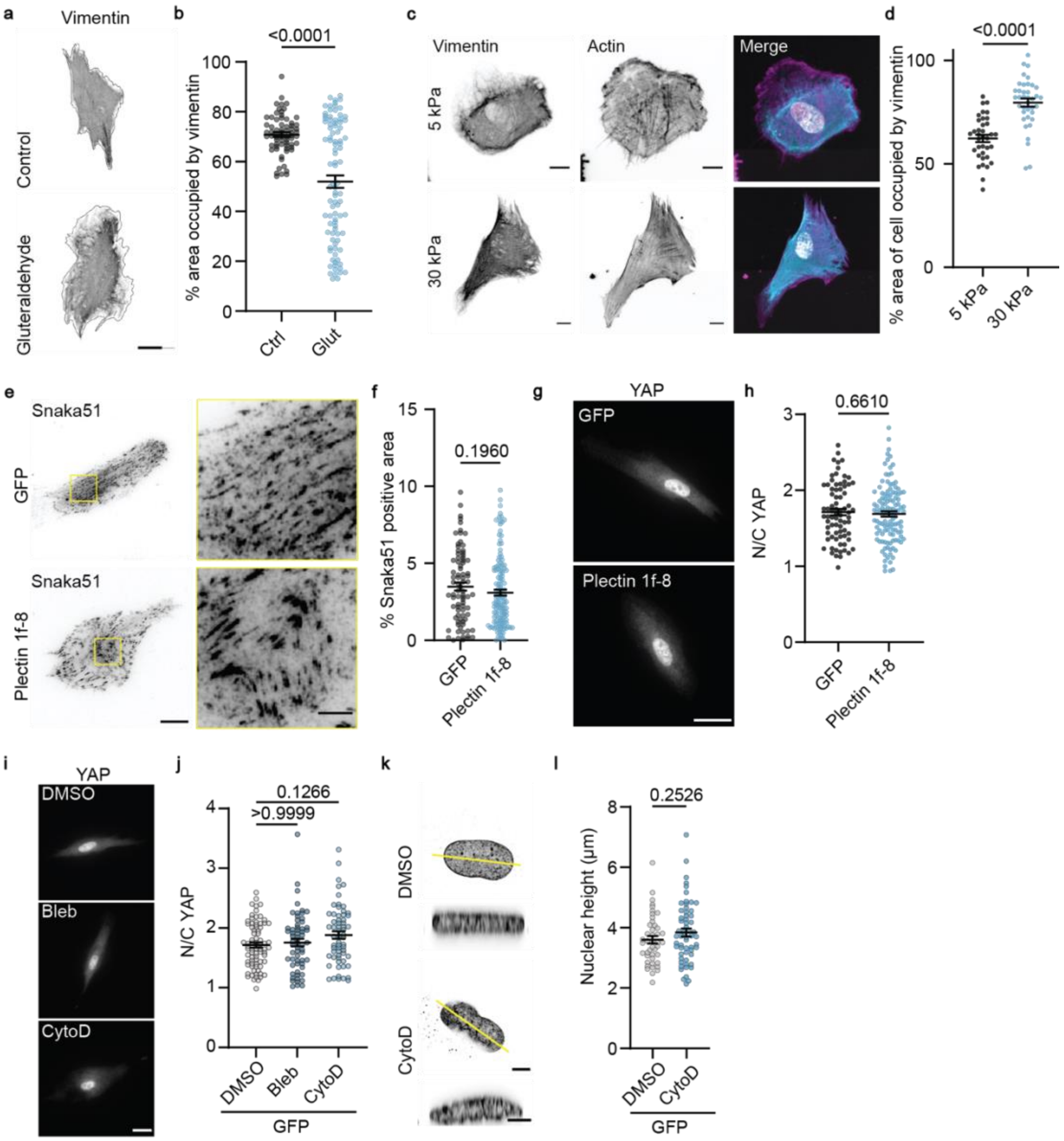


**Supplementary figure 4. Additional characterization of the relationship between fibrillar adhesion formation and vimentin morphology. a.** Example images of vimentin stained cells on control or glutaraldehyde blocked substrates; scale bar 25µm. **b.** Quantification of the percentage area of the cell occupied by vimentin on cells seeded on control or glutaraldehyde blocked substrates. (n=68/93 cells for control/glut from 5 independent experiments; Mann-Whitney test). **c.** Confocal images of cells seeded on 5 kPa and 30 kPa polyacrylamide gels stained with vimentin, actin (phalloidin) and the nucleus (hoechst). All images are a z-projection (sum) of 21 confocal slices. Scale bars are 10 µm. **d.** Quantification of the percentage area of the cell occupied by the vimentin network on cells seeded on 5kPa gels and 30kPa gels. (n=36/38 cells for 5kPa/30kPa from 3 independent experiments; Mann-Whitney test). **e.** Example images of snaka51 stained cells transfected with GFP or plectin 1f-8-GFP; scale bar 25µm/ zoom 5µm. **f.** Quantification of percentage area under the nucleus occupied by fibrillar adhesions for cells transfected with GFP control of plectin 1f-8-GFP mutant. (n=83/134 cells from 2 independent experiments; Mann-Whitney test). **g.** Example images of YAP staining for cells expressing GFP or plectin 1f-8-GFP. Scale bar 25 µm. **h.** Quantification of N/C YAP ratio for cells transfected with GFP and plectin 1f-8-GFP. (GFP n=76, plectin 1f-8-GFP n=120, from at least 3 independent experiments; unpaired t-test). **i.** Control GFP transfected cells subjected to 30 minute pharmacological treatment stained with YAP antibody- Scale bar 25µm. **j.** Analysis of N/C YAP after 30 minute pharmacological treatment to GFP transfected cells. (n=76/61/62 cells for DMSO/bleb/cytoD from 3 independent experiments; Kruskal-Wallis test with Dunn's multiple comparison test). **k.** LaminB stained nuclei of GFP transfected cells treated with DMSO or CytoD. Scale bar 5µm. **l.** Quantification of nuclear height for GFP transfected cells treated with vehicle DMSO or CytoD for 1 hour. (n=44/58 cells for DMSO/cytoD from 3 independent experiments; Mann-Whitney test).

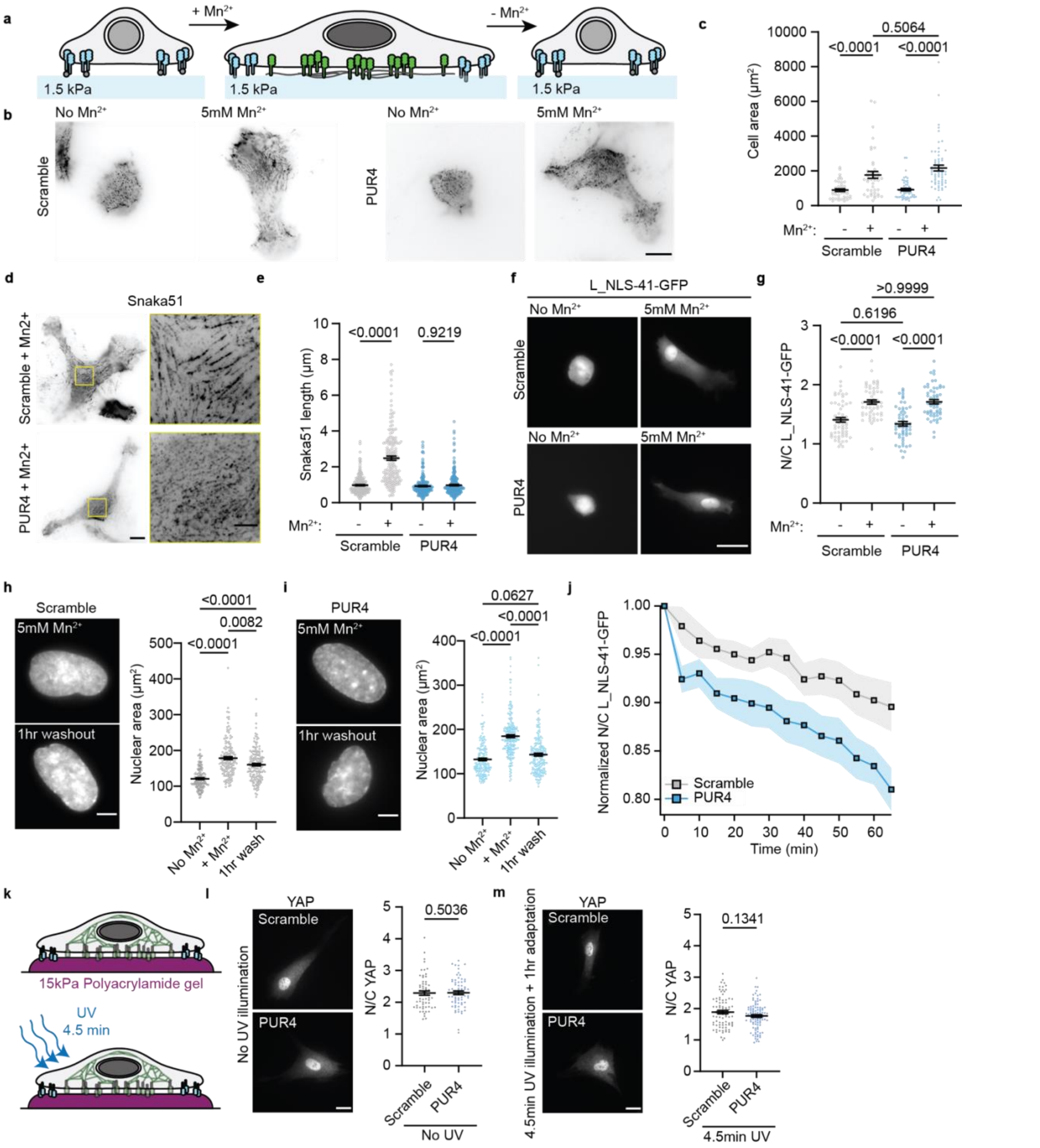


**Supplementary figure 5. Additional characterization of cellular adaptation timescales depending on fibrillar adhesion formation. a.** Schematic representation of the experimental design where cells were seeded on soft (1.5 kPa) fibronectin coated polyacrylamide gels. Cells were then incubated with $Mn^{2+}$ to trigger the activation of integrin α5β1, leading to cell spreading and a mechanically active phenotype. Finally, the $Mn^{2+}$ was washed out, and the cells adaptation time was monitored. **b.** Example images of Snaka51 stained cells seeded on 1.5kPa gels in the absence or presence of 5mM $Mn^{2+}$, in scramble and PUR4 containing media. Scale bar 20µm. **c.** Quantification of cell area for scramble and PUR4 in the absence or presence of $Mn^{2+}$. (Scramble n=43, scramble with $Mn^{2+}$ n=54, PUR4 n=57, PUR4 with $Mn^{2+}$ n=64 cells, from 3 independent experiments; 2-way ANOVA with Tukey's multiple comparison test). **d.** Example images of snaka51 stained cells treated with $Mn^{2+}$ in scramble or PUR4 peptide. Scale bar 20µm / 5µm zoom. **e.** The addition of $Mn^{2+}$ to cells in scramble peptide trigger a significant increase in fibrillar adhesion length, but do not in the presence of the PUR4 peptide. (Scramble n=240 adhesions (48 cells), Scramble with $Mn^{2+}$ n=170 (34), PUR4 n=230 (46), PUR4 with $Mn^{2+}$ n=280 (56); from 3 independent experiments; 2-way ANOVA with Tukey's multiple comparison test). **f.** Example images of L_NLS-41-GFP transfected cells seeded on 1.5kPa gels in absence or presence of $Mn^{2+}$ for scramble and PUR4 peptide. Scale bar 20µm. **g.** Quantification of the N/C L_NLS-41-GFP ratio in absence or presence of $Mn^{2+}$ for cells in scramble or PUR4 peptide. (Scramble n=61, scramble with $Mn^{2+}$ n=55, PUR4 n=57, PUR4 with $Mn^{2+}$ n=55 cells from 3 independent experiments; 2-way ANOVA with Tukey's multiple comparison test). **h.** Nuclear area of cells seeded on 1.5kPa gels in the absence, presence or after 1 hour washout of $Mn^{2+}$ activation in the scramble peptide. The nuclear area after 1 hour washout is still significantly larger than in the absence of $Mn^{2+}$ activation. (n=157/151/160 cells for No-$Mn^{2+}$/With- $Mn^{2+}$/ $Mn^{2+}$-washout from 5 independent experiments; Kruskal-Wallis test with Dunn's multiple comparisons test). **i.** Nuclear area of cells seeded on 1.5kPa gels in the absence, presence or after 1 hour washout of $Mn^{2+}$ activation in the PUR4 peptide. The nuclear area after 1 hour washout is not significantly different than in the absence of $Mn^{2+}$ activation. (n=164/203/192 cells for No-$Mn^{2+}$/With- $Mn^{2+}$/ $Mn^{2+}$-washout from 5 independent experiments; Kruskal-Wallis test with Dunn's multiple comparisons test). **j.** The change in nuclear/cyto ratio of L_NLS-41-GFP (normalized to pre-washout value) upon removal of the $Mn^{2+}$ activation for scramble (grey) and PUR4 (blue) condition. Cells were seeded on 1.5kPa gels, activated with $Mn^{2+}$ for a minimum of 4 hours prior to washout. Scramble n=37, PUR4 n=35 from 3 independent experiments. **k.** Schematic of experimental set-up whereby cells are seeded on 15kPa polyacrylamide gels (PAA) that do not change mechanical properties upon UV illumination. Cells were seeded for a minimum of 4 hours prior to 4.5min UV illumination in the presence of the scramble or PUR4 peptide. Cells were subsequently left for 1 hour to adapt and then fixed and stained. **l.** Immunostaining of YAP in cells seeded on 15kPa PAA gels without UV exposure, cultured in scramble or PUR4 peptide. Scale bar 20µm. Quantification of N/C YAP ratio for scramble or PUR4 conditions prior to UV illumination. Mann-Whitney test from 2 independent experiments, Scramble n=60 cells, PUR4 n=68 cells. **m.** Immunostaining of YAP in cells seeded on 15kPa PAA gels with 4.5min *in-situ* UV exposure followed by 1 hour adaptation time, cultured in scramble or PUR4 peptide. Scale bar 20µm. Quantification of N/C YAP ratio for scramble or PUR4 conditions subjected to 4.5min UV illumination and left for 1 hour. Mann-Whitney test from 2 independent experiments, Scramble n=83 cells, PUR4 n=98 cells.

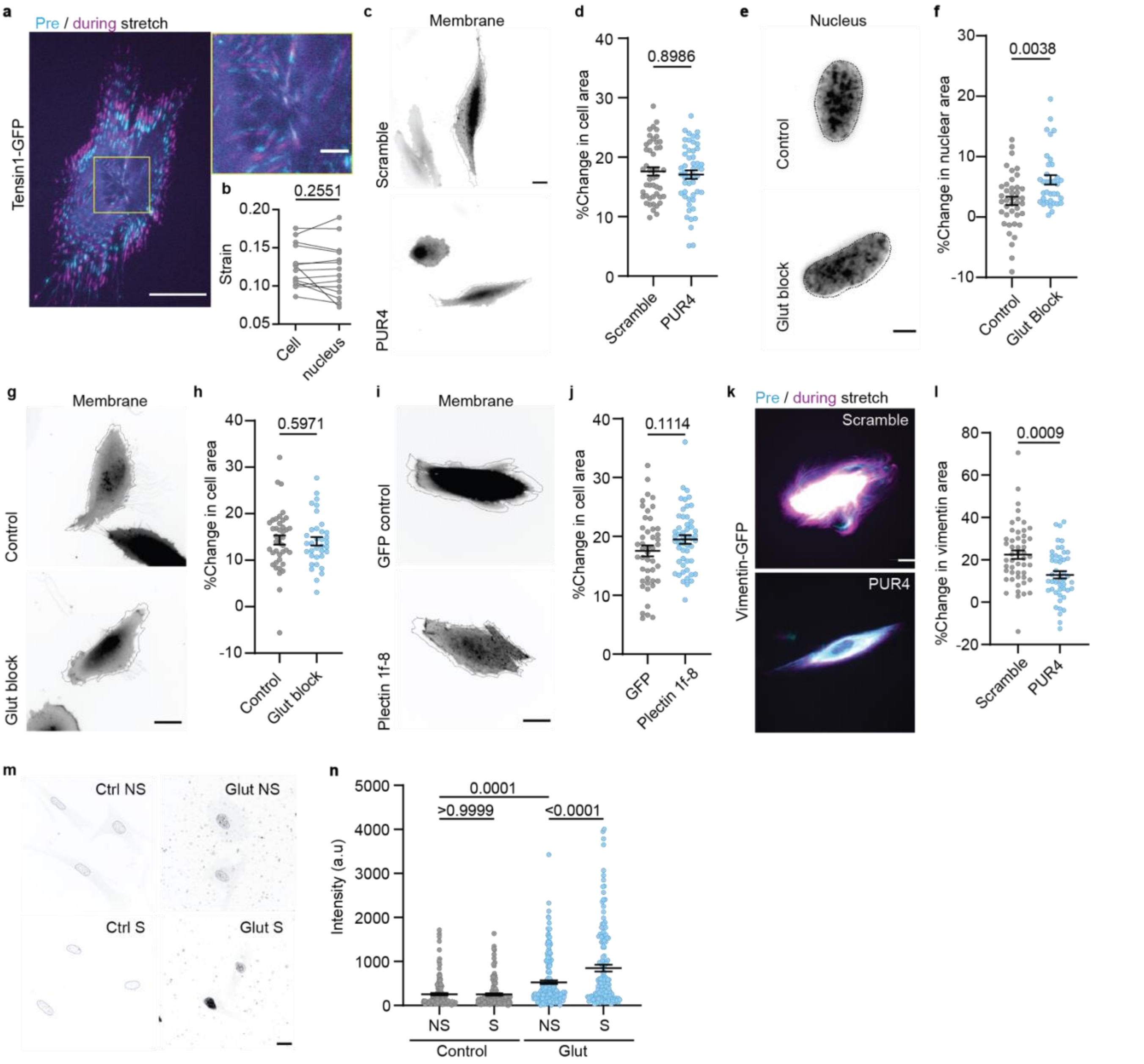


**Supplementary figure 6. Additional characterization of the effect of stretch depending on the existence of fibillar adhesions. a.** Image of a tensin1-GFP transfected cell before (cyan) and during (magenta) stretch. Pre and during stretch images are aligned at an adhesion located centrally under the nucleus. Scale bar is 25 µm and 5 µm on zoomed inset. **b.** Analysis of the whole cell strain and the strain under the nucleus (analysed from the change in distance between two tensin1-GFP adhesions located under the nucleus) upon a 10% stretch. There is no significant difference between the whole cell strain and subcellular strain. P-values calculated by a paired two-tailed t-test from 3 independent experiments (n=14 cells). **c.** Cell membrane before and during (dashed line) stretch for cells cultured in scrambled or PUR4 peptide. Scale bar 20µm. **d.** Quantification of the percentage change of cell area between the scramble and PUR4 condition. (Scramble n=47, PUR4 n=50 cells from 3 independent experiments; Mann-Whitney test). **e.** Images of Hoechst stained nuclei before (image) and during (dashed line) stretch for control cells, and cell seeded on gluteraldehyde blocked PDMS substrates. Scale bar 5µm. **f.** The percentage change in nuclear area upon stretch for control and gluteraldehyde cells are statistically significant. (Control n=41, cells glut n=35 cells, from 3 independent experiments; Mann-Whitney test). **g.** Images of the cell membrane before and during (dashed line) the application of stretch for control cells and cells seeded on gluteraldehyde blocked substrates. Scale bar 20µm. **h.** Quantification of the percentage change of cell area for cells seeded on control or gluteraldehyde blocked substrates. (control n=41 cells, glut n=35 cells from 3 independent experiments; Mann-Whitney test). **i.** Images of cells transfected with GFP-only or Plectin 1f-8-GFP before and during stretch. Scale bar 20µm. **j.** Quantification of the percentage change in cell area between GFP-only or Plectin 1f-8-GFP cells. (GFP n=46 cells, Plectin 1f-8 n=51 cells from 3 independent experiments; Mann-whitney test). **k.** Images of vimentin-GFP transfected cells in scramble or PUR4 peptide before (cyan) and during (magenta) mechanical stretch. Scale bar is 20 µm. **l.** Quantification of the percentage change in the vimentin area upon stretch for the cells in the scramble peptide compared to the PUR4 peptide. (Scramble n=50 cells, PUR4 n=47 cells from 3 independent experiments. Mann-Whitney t-test). **m.** Immunostaining of γH2Ax in control or glut blocked cells that were not stretched (NS) or subjected to stretch (S). Scale bar 20µm. **n.** Nuclear intensity of γH2Ax marker. In control cells there is no significant difference between NS and S conditions. In the glut block condition, stretch triggers a significant increase in the nuclear intensity of γH2Ax. (Control NS n=156, control S n=138, glut NS n=196, glut S n=140 nuclei from 2 independent experiments; 2-way ANOVA with Tukey's multiple comparison test).

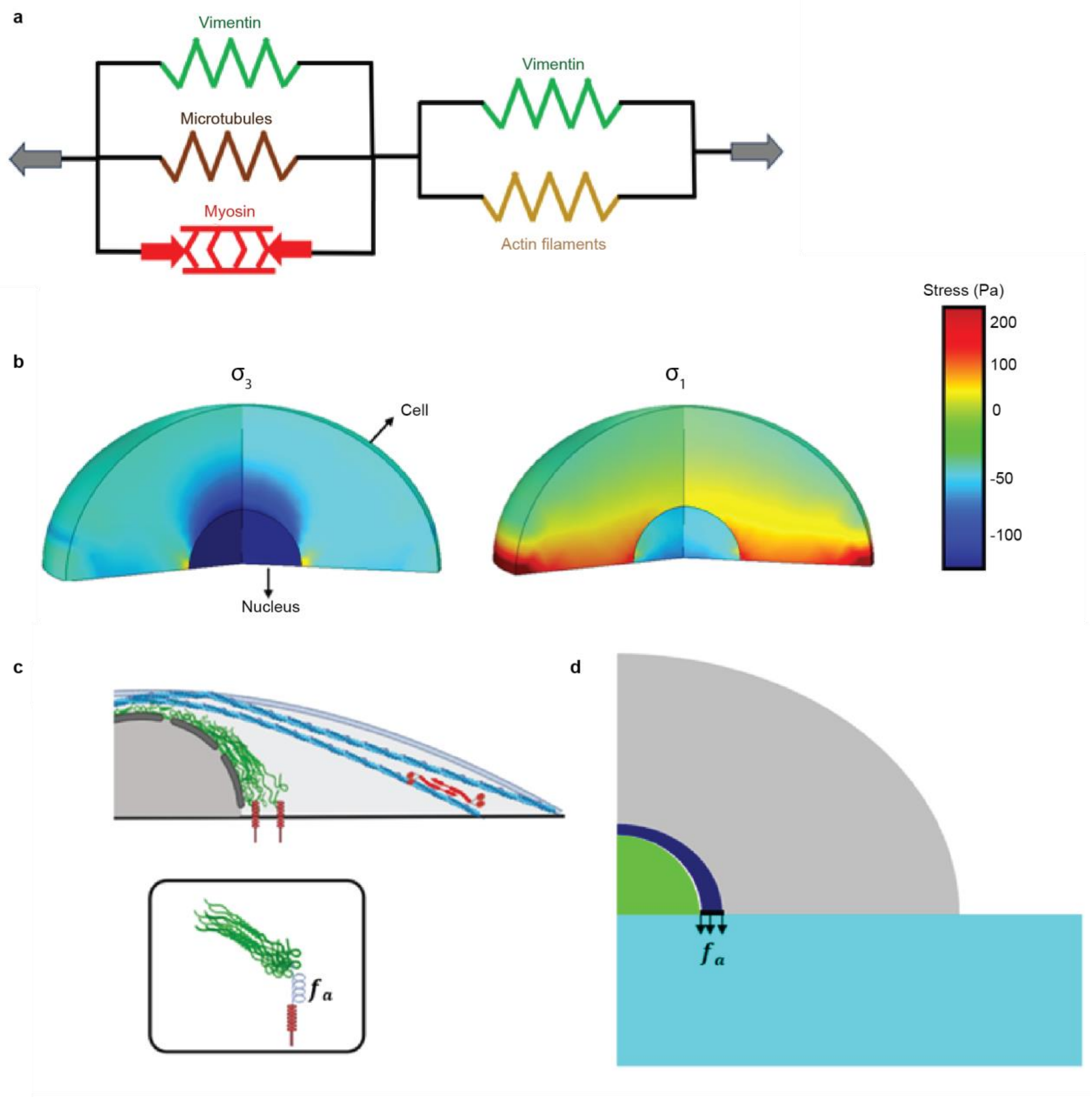


**Supplementary Figure 7. Additional characterisation of the computationally modelling methods. a.** A one dimentional representation of the chief components in the computational model. Vimentin intermediate filaments in direct physical contact with actin are in tension and essential for force transmission to the ECM while VIFs adjacent to the nucleus reinforce the microtubules and are under compression. **b.** Principal Stress distribution within the cell due to contractile forces, where $\sigma_1 > \sigma_2 > \sigma_3$. Maximum compressive stress field is found around the nucleus and represents the formation of vimentin cage, while the maximum tensile stress is observed along the basal plane and close to the cell periphery. **c.** Schematic representation of the engagement of vimentin cage with fibrillar adhesions due to cell contraction. **d.** Schematic of model geometry describing how the adhesive forces are applied to represent anchoring by adhesions.
